# Mapping Projectome Heterogeneity of Subiculum Neuron Cell Types

**DOI:** 10.64898/2026.04.01.716004

**Authors:** Ann W. Saustad, Michael S. Bienkowski

## Abstract

The subiculum (SUB) is the main output structure of the hippocampus, influencing diverse behaviors through its widespread cortical and subcortical connections. Our previous work creating the mouse Hippocampus Gene Expression Atlas (HGEA) identified four genetically distinct cellular layers across five columnar domains in the SUB, with gene expression boundaries corresponding to distinct connectivity patterns and brain-wide networks involved in spatial navigation, social behavior, and neuroendocrine regulation (Bienkowski et al., 2018). Using the Digital Brain Mouse Projectome Atlas (MPA) tool, we conducted ‘virtual tract-tracing’ to assess whether connectivity patterns of single-neuron 3D reconstructions aligned with HGEA-defined SUB cell types (Qiu et al., 2024). We classified 689 SUB projection neurons into 12 HGEA cell-type groups based on their laminar and columnar distributions, whose spatial organization recapitulated HGEA-defined 3D boundaries. Using this population sample, we performed a SUB cell-type census, characterized neuronal heterogeneity and projection prevalence, identified common and rare connectivity motifs and axonal collateralization patterns, and defined distinct projection themes for each SUB cell type. Together, this analysis integrates single-neuron and population-level data to advance understanding of SUB cell type organization and its contributions to brain-wide networks regulating diverse behaviors.

## I: Introduction

The subiculum (SUB) is an allocortical brain structure that is anatomically positioned as a major gateway for hippocampal output and influencing a diverse array of animal behaviors. In rodents, the SUB is classically divided into dorsal and ventral subregions (SUBd and SUBv, respectively) based on its bisection by the caudal CA1 in coronal sections and lesions of these regions produce functional deficits in different behaviors (Fanselow & Dong, 2010; Moser & Moser, 1998). However, gene expression and anatomical tracing studies spanning the last several decades have revealed that the anatomical organization of the SUB is more complex, with both columnar and laminar organization of neuronal cell types (Bienkowski et al., 2018; Boccara et al., 2015; Strange et al., 2014; Kishi et al., 2000).

In our previous study creating the Hippocampus Gene Expression Atlas (HGEA), we integrated gene expression and connectivity data to identify four genetically-distinct layers of cell types distributed across five columnar regions constituting the entire mouse SUB (Bienkowski et al., 2018). Subsequent gene expression analysis in humans suggest that this columnar and laminar organization is conserved across mammalian species, albeit with divergent combinatorial gene expression patterns (Bienkowski et al., 2021). Recently, high-throughput single neuron projectome studies, such as the Mouse Projectome Atlas (MPA) and the Janelia MouseLight Project, have provided unprecedented detail of single-cell connectivity across the whole mouse brain (Qiu et al., 2024; Winnubst et al., 2019). These datasets offer a new opportunity to evaluate how SUB cell type connectivity at the single-cell level maps onto the HGEA columnar and laminar framework, and to analyze connectivity-based cell type heterogeneity and axon collateralization patterns that are critical for understanding systems-level information processing. Using the Mouse Projectome Atlas Digital Brain (www.digital-brain.cn), we performed virtual tract tracing experiments on 689 3D-reconstructed MPA SUB neurons and classified each neuron into an HGEA-defined cell type based on its soma location and projection pattern. Here, we present the results of this classification, characterize the connectivity motifs and collateralization patterns that define each SUB cell type, and discuss their implications for brain-wide networks regulating navigation, memory, and affect.

## II: Subiculum

### a. Neuroanatomical Organization of the Subiculum

Early anterograde and retrograde tracing studies provided initial evidence for both a columnar and laminar organization of SUB pyramidal cell types. Anterograde injections along the longitudinal axis of the SUB revealed a topographic organization of the SUB axonal projections to the septum and hypothalamus (Swanson & Cowan, 1975). Rather than forming a continuous gradient, injections targeting specific domains within the SUB labeled distinct, but partially overlapping topographic subregions in these target areas. In rats, retrograde tracer injections into brain areas innervated by the SUB revealed laminar distributions of labeled neurons within the SUB pyramidal layer (Ishizuka, 2001). Although a sublayer organization was clearly present within the SUB pyramidal layer, retrograde labeling only partially captured the full laminar organization, as tracer injection sites cannot typically encompass an entire target region while maintaining anatomical specificity. As a result, tracing experiments inherently subsample connectivity pathways, making it difficult to determine the total number of layers and their full distribution across the SUB axis (Saleeba et al., 2019).

Complementing the anatomical retrograde labeling patterns, laminar gene expression patterns within the SUB were later identified using *in situ* hybridization data from the Allen Brain Atlas database (Lein et al., 2007). Genes associated with inhibitory neurons are expressed broadly and sparsely across both the SUB pyramidal and molecular layers. In contrast, genes associated with excitatory neurons and other neurochemical systems are expressed primarily within the pyramidal layer, with combinatorial laminar patterns extending across the entire SUB axis. By analyzing these combinatorial expression patterns in creating the HGEA, we identified four neuronal sublayers within the SUB pyramidal layer that are differentially distributed along the longitudinal axis (SUB_1, SUB_2, SUB_3, SUB_4; (Bienkowski et al., 2018); **Fig. 1A**). Briefly, SUB_1 occupies a superficial position within the pyramidal layer of the distal dorsal SUB and continues to the caudal end of the SUB, where it curves ventrally and slightly back rostrally, forming a C-shaped trajectory. SUB_2 contains the most superficial neurons of the ventral SUB, with SUB_3 occupying the layer immediately deep to SUB_2. At the caudal levels where CA1 ends, both layers extend dorsally to become adjacent to SUB_1 in the distal dorsal SUB. Here, SUB_3 but not SUB_2, continues rostrally adjacent to SUB_1, forming the superficial layer of the ProSUB. SUB_4 is the deepest layer across all SUB regions, positioned closest to the alveus white matter, and likely corresponds to the polymorphic cell layer described by Cajal, which contains pyramidal, stellate, triangular, and fusiform neurons (DeFelipe & Fariñas, 1992; Ramón y Cajal, 1909). SUB_4 is present across all parts of the SUB and is generally thin (1-2 cells wide) but thickens at both the dorsal and ventral tips, forming characteristic bulb-shaped endings. In three-dimensions, these cell type layers resemble sheets or tectonic plates that have shifted atop or beneath each other across the hippocampal axis. How these layers are represented in specific parts of the SUB is an important distinction for how regional areas of the SUB form structural/functional columns for processing hippocampal inputs and distributing encoded neural signals to downstream brain networks.

**Figure 1.**
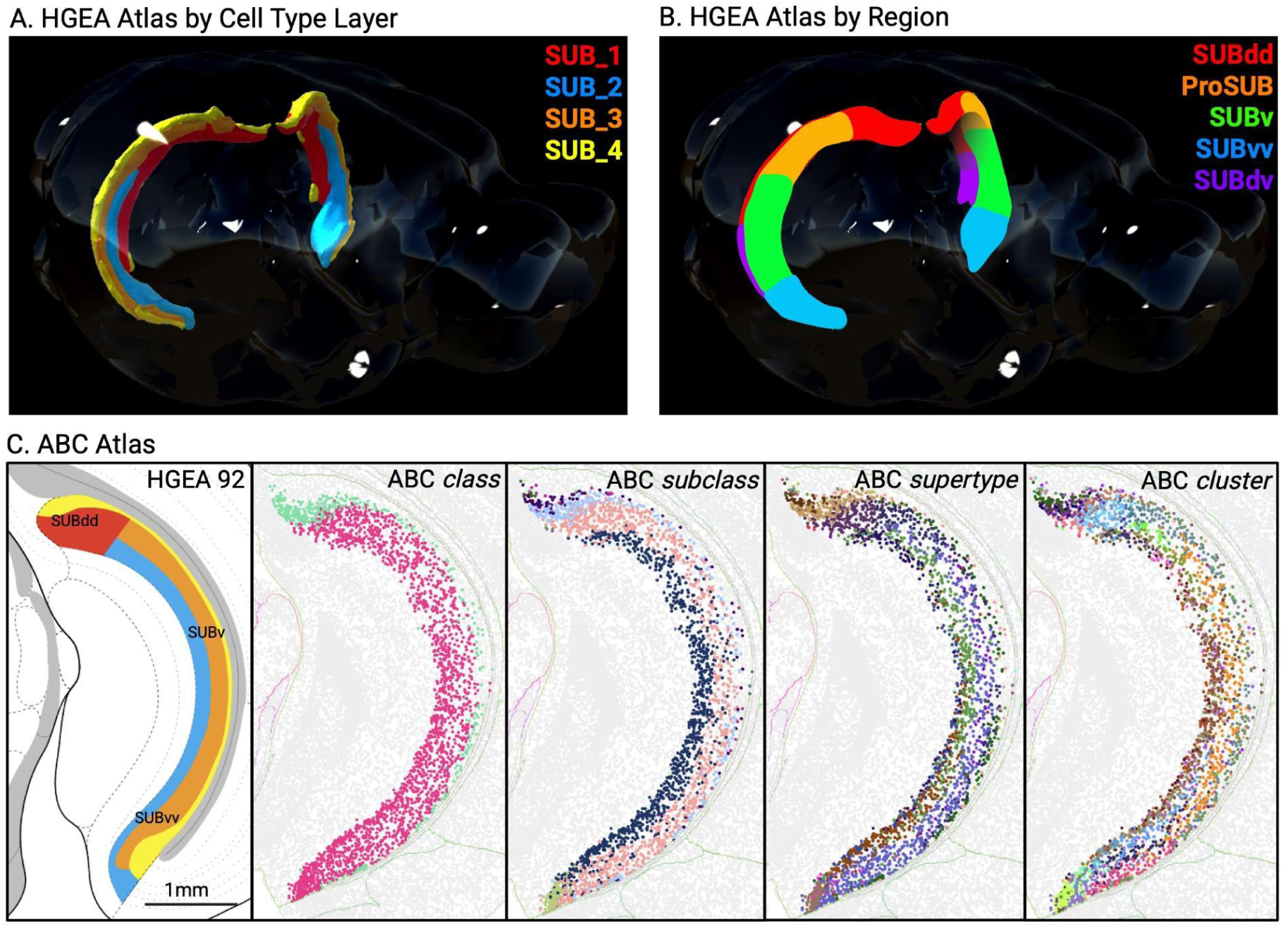
HGEA and Allen Brain Cell (ABC) atlases reveal laminar and regional organization of subicular neurons. **(A)** 3D rendering of HGEA-defined subiculum cell-type layers (SUB_1–SUB_4). **(B)** 3D rendering of HGEA-defined subicular regions, including SUBdd, ProSUB, SUBv, SUBvv, and SUBdv. **(C)** Schematic representation of laminar and regional boundaries at HGEA level 92, with color coding corresponding to SUB_1 (red), SUB_2 (blue), SUB_3 (orange), and SUB_4 (yellow) in panel 1. MERFISH spatial transcriptomic data from the Allen Brain Cell (ABC) atlas illustrating the hierarchical classification of subicular neurons across four progressively granular annotation levels: class, subclass, supertype, and cluster (left to right). Scale bar, 1 mm.

### b. Integrating Gene Expression and Connectivity to Understand Laminar and Columnar Organization

To validate that gene expression boundaries correspond to differences in anatomical connectivity, we systematically compared HGEA gene expression patterns with retrograde tracing data from the Mouse Connectome Project. This integration revealed that each of the four SUB pyramidal layers exhibits relatively distinct projection patterns, confirming that combinatorial gene expression delineates anatomically and functionally distinct cell populations (Bienkowski et al., 2021).

Just as variation in laminar composition defines distinct cortical areas with specialized connectivity (Brodmann areas; (García-Cabezas et al., 2019), the differential distribution of SUB layers along the dorsoventral axis creates five columnar domains (**Fig. 1B**). The distal SUB is bilaminar, containing layers 1 and 4, and extends along a curved trajectory from its rostral-dorsal position (SUBdd), corresponding to the classical dorsal subiculum, to a caudal-ventral extension (SUBdv), where layer 4 forms the characteristic bulb-shaped enlargements at the dorsal and ventral tips. The prosubiculum (ProSUB), located adjacent to CA1, is also bilaminar but composed of layers 3 and 4, consistent with the position and cellular composition of Lorente de Nó’s original description (Lorente De Nó, 1934; Rosenblum et al., 2024). The ventral subiculum (SUBv) has a trilaminar organization with layers 2, 3, and 4, while its most ventral portion (SUBvv) exhibits altered layer proportions and connectivity patterns more closely resembling ProSUB than SUBv proper.

This laminar-columnar organization reveals that the classical division of the SUB into dorsal and ventral regions, based on their separation by CA1 at rostral levels, does not accurately reflect the underlying cellular architecture. At caudal levels where CA1 ends, these classically defined regions become continuous, with layers 2 and 3 extending dorsally to merge with layer 1 territories. The SUB is therefore better understood as two major divisions with fundamentally different laminar architectures: SUBdd/SUBdv (layers 1 and 4) versus ProSUB/SUBv/SUBvv (layers 2, 3, and 4; layer 2 absent in ProSUB). These divisions exhibit distinct brain-wide connectivity patterns, suggesting they serve fundamentally different functional roles in transforming hippocampal output.

While the HGEA is based on both genetic and connectomic data, subsequent single-cell RNA sequencing studies have validated and extended these gene expression patterns. Cembrowski et al. (2018) identified structural and functional differences between proximal and distal dorsal SUB, corresponding to ProSUB and SUBdd, respectively, and delineated 8 pyramidal cell subclasses with discrete laminar and columnar spatial domains (Cembrowski, Phillips, et al., 2018; Cembrowski, Wang, et al., 2018). Alternative nomenclatures have since been proposed (Ding et al., 2020), though the precise correspondence between these frameworks and the HGEA organization remains to be fully resolved. More recently, the Allen Brain Cell (ABC) Atlas integrated single-cell and spatial transcriptomics across the entire mouse brain, providing high-resolution molecular definitions of SUB cell types organized into hierarchical levels: class, subclass, supertype, and cluster (Yao et al., 2023). The HGEA laminar organization aligns with intermediate levels of this hierarchy, with SUB_2 corresponding to the CA1-ProS subclass, SUB_1 and SUB_3 to SUB-ProS glutamatergic supertypes, and SUB_4 encompassing a combination of deep-layer subclasses including corticothalamic and intratelencephalic glutamatergic neurons (**Fig. 1C**).

### c. Single-Cell Projectomes Enable Analysis of Projection Neuron Diversity

Classical anatomical studies using retrograde and anterograde tracing suggested that individual SUB neurons exhibit relatively limited axonal collateralization, with most neurons projecting to single target regions (O’Mara et al., 2001; Witter, 2006). This view, which contrasted SUB organization with the extensive collateralization observed in hippocampal CA fields (Swanson, 1981), was reinforced by the observation that SUB projections are organized in a largely columnar and laminar fashion, with distinct neuronal populations innervating different downstream targets (Ishizuka, 2001; Witter, 2006). More recent double-labeling studies have demonstrated that SUB collateralization is more prevalent than previously appreciated, with approximately half of dorsal SUB neurons projecting to the retrosplenial cortex also innervating the mammillary bodies, and additional populations targeting both the mammillary body and entorhinal cortex (Aggleton & Christiansen, 2015; Kinnavane et al., 2018). These findings indicate that the extent of SUB axonal collateralization was substantially underestimated by earlier single-label tracing studies.

Recent advances in sparse viral labeling combined with whole-brain imaging have enabled comprehensive reconstruction of individual neuron morphologies at unprecedented resolution. The Janelia MouseLight project pioneered this approach, using two-photon tomography to reconstruct over 1,000 complete projection neurons across the mouse brain (Winnubst et al., 2019). Building on this methodology, the CEBSIT(ION) Digital Brain Mouse Projectome Atlas (MPA) completed digital reconstruction of over 10,000 individual neurons in the hippocampus and subicular complex using fluorescence micro-optical sectioning tomography (fMOST; (Qiu et al., 2024)). This openly accessible dataset (https://mouse.digital-brain.cn/hipp) provides complete 3D axonal morphologies at single-cell resolution, enabling systematic re-evaluation of SUB collateralization patterns and projection diversity across the entire HGEA laminar-columnar framework.

Single-cell projectome studies in the neocortex have demonstrated that projection neurons can be classified into distinct collateralization types based on their axonal targets: intratelencephalic (IT) neurons projecting bilaterally to cortex and striatum, extratelencephalic (ET) neurons with ipsilateral descending subcortical projections, and corticothalamic (CT) neurons targeting the thalamus (Mao & Staiger, 2024; Harris et al., 2019; Winnubst et al., 2019). Population-level tracing data from the HGEA suggested that SUB layers exhibit projection patterns analogous to these cortical types, with SUB_1 and SUB_3 resembling layer 5 ET neurons, SUB_2 exhibiting layer 2/3-like IT connectivity, and SUB_4 showing mixed CT/IT patterns characteristic of layer 6 (Bienkowski et al., 2018). Whether individual SUB neurons conform to these broad classes, and whether their collateralization patterns are organized systematically across HGEA layers, has not been fully resolved at the single-cell level.

## III: Methodology and Results

### a. Virtual Tract Tracing to Identify and Classify HGEA SUB Cell Types at the Single-Cell Level

To systematically characterize SUB projection patterns at single-cell resolution and test whether individual neurons conform to HGEA layer-specific connectivity profiles, we performed virtual tract tracing on neurons from the Mouse Projectome Atlas (MPA). The MPA Digital Brain (https://mouse.digital-brain.cn/projectome) currently contains over 45,000 publicly available single-neuron reconstructions registered to the Allen CCFv3 reference atlas. Since the MPA dataset uses CCFv3 anatomical definitions, HGEA SUB cell types were expected to distribute across CCFv3 subiculum (SUB), prosubiculum (ProS), and potentially the hippocampo-amygdalar transition area (HATA) subregions. According to Ding et al. (2020), who established the CCFv3 SUB nomenclature, the CCFv3 SUB region should correspond primarily to HGEA SUBdd/SUBdv (containing layers 1 and 4), while CCFv3 ProS should align with HGEA ProSUB (containing layers 3 and 4).

Using retrograde labeling patterns that were found to relatively specifically label the SUB cell type layers in the HGEA (Bienkowski et al., 2018), we queried the MPA database using soma location and projection target filters characteristic of each layer: SUB_1 neurons projecting to ventral retrosplenial cortex (RSPv), SUB_2 neurons projecting to taenia tecta (TT), SUB_3 neurons projecting to nucleus accumbens (ACB), and SUB_4 neurons projecting to nucleus reuniens (RE). We initially restricted our search to neurons with somata in the CCFv3 SUB region, expecting to identify primarily SUB_1 and SUB_4 cell types consistent with the proposed correspondence to HGEA SUBdd/SUBdv correspondence. Instead, we identified neurons with projection patterns characteristic of all four HGEA layers within the CCFv3 SUB boundaries, indicating that the CCFv3 anatomical delineations do not precisely align with the HGEA gene expression-based framework. This discrepancy between anatomical atlases has important implications for interpreting connectivity data across studies using different coordinate systems (see Discussion).

To comprehensively characterize SUB projection diversity, we expanded our analysis to include the CCFv3 HATA region and performed rigorous manual curation of all identified neurons. Given that MPA reconstructions are registered from multiple individual brains to a 10μm isotropic resolution atlas, some spatial imprecision is expected. We manually verified soma locations and axonal trajectories for 689 neurons and classified each into HGEA cell types based on combined soma position and projection pattern criteria (**Table 1**). When visualized in 3D space, the color-coded MPA reconstructions recapitulated the laminar and columnar organization defined by the HGEA (**Fig. 2A,B**), with clear topographic axonal connectivity patterns evident for both laminar and columnar groupings (**Fig. 2C,D**). For each HGEA region and layer, we quantified the proportion of neurons innervating 23 major brain structures (**Table 2**) and generated radar plots to visualize the differential connectivity profiles of each SUB region (**Fig. 3**) and cell-type layer (**Fig. 4**). Together, these analyses enabled us to evaluate individual neuronal conformity to layer-specific patterns and to characterize the extent of axonal collateralization across functionally distinct brain networks.

**Figure 2.**
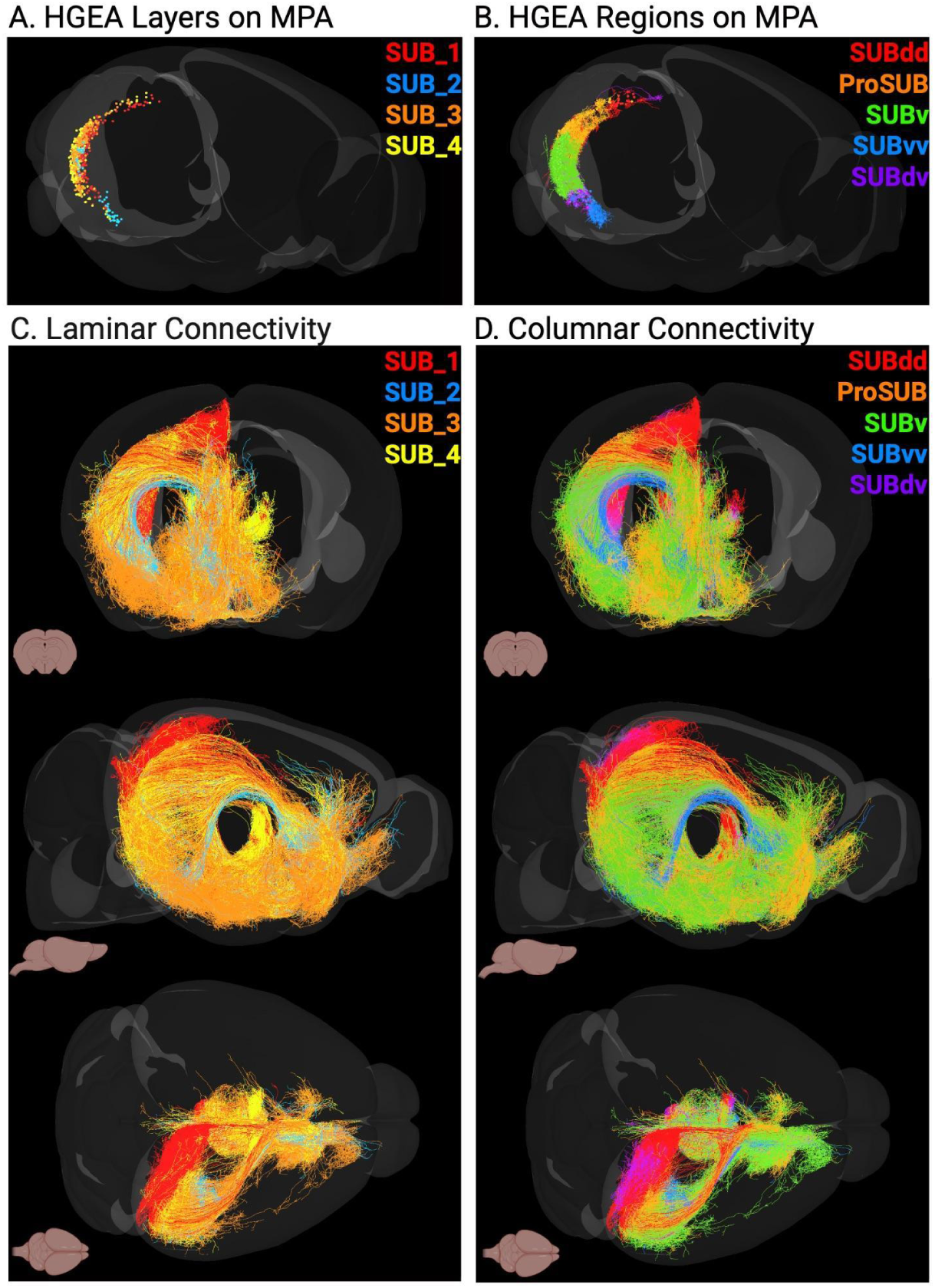
HGEA laminar and columnar organization of MPA subiculum neuron reconstructions. **(A)** Somas of 689 MPA SUB neurons classified by HGEA layers (SUB_1-4) and **(B)** Somas and dendrites of the same 689 MPA SUB neurons organized by HGEA regions (SUBdd, ProSUB, SUBv, SUBvv, and SUBdv). Reconstructions of 689 MPA SUB neuron soma, dendrites, and axons classified by their laminar **(C)** or columnar **(D)** organization exhibit distinct topographical connectivity patterns across the mouse brain.

**Figure 3.**
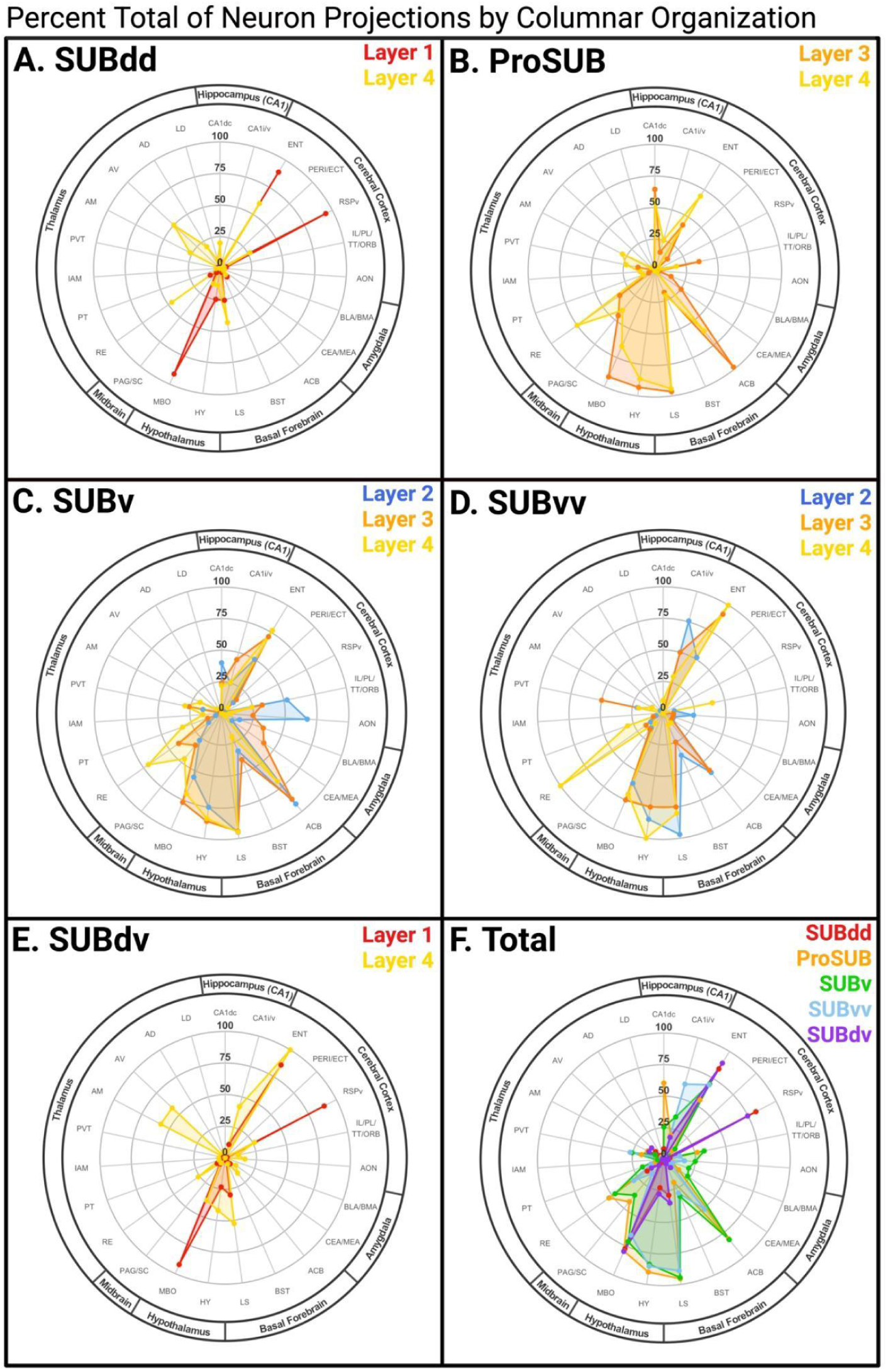
Radar plots of the percent total of subicular projections from HGEA-defined columns across brain regions. Radar plots showing percent total projections from each HGEA-defined column, separated by laminar identity. **(A)** SUBdd, **(B)** ProSUB, **(C)** SUBv, **(D)** SUBvv, **(E)** SUBdv, and **(F)** Total SUB Columns. Axes represent target brain regions; values indicate percentage of total projections for each columnar and laminar group (Table 2).

**Figure 4.**
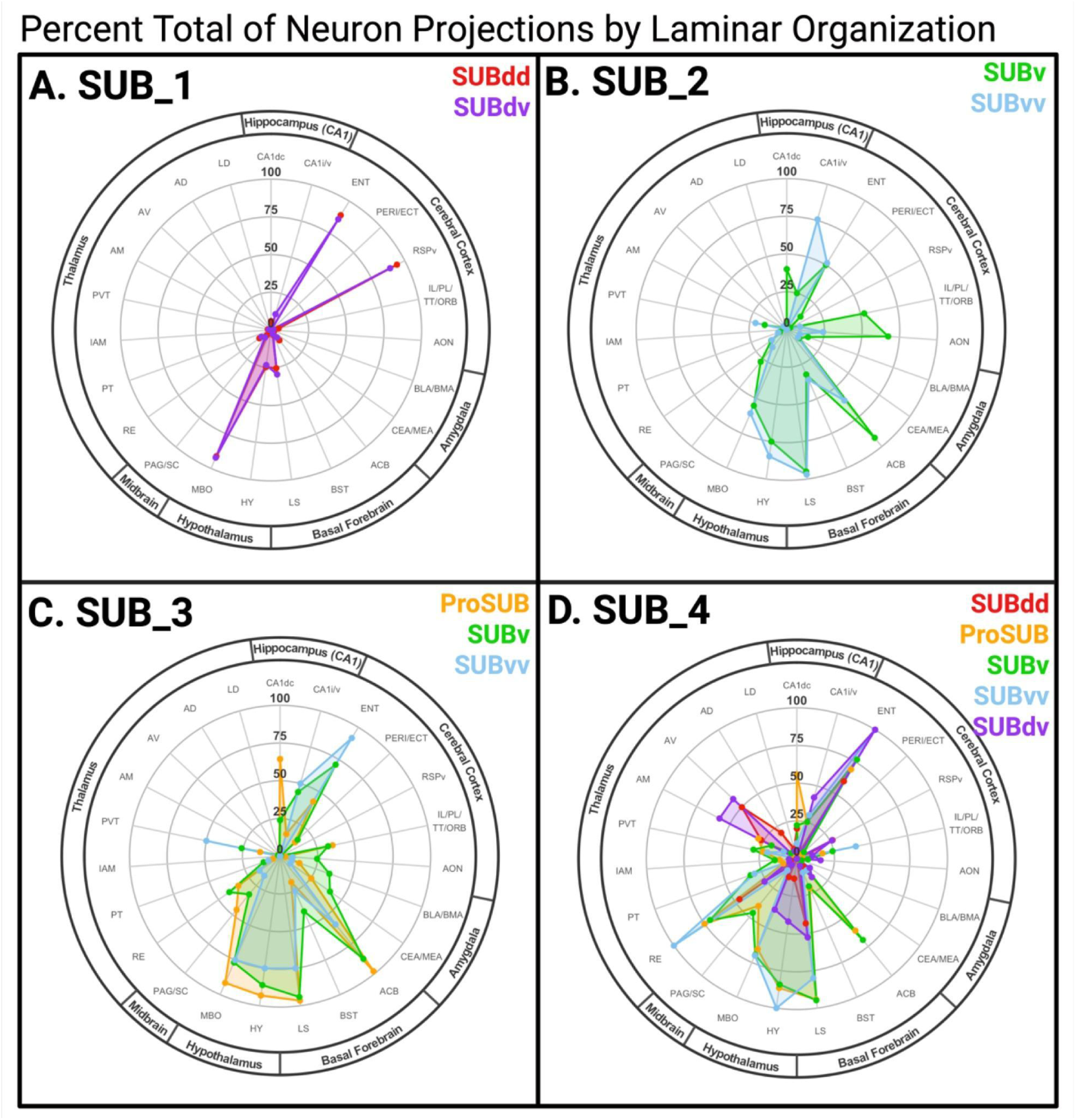
Radar plots of the percent total of subicular projections from HGEA-defined layers across brain regions. Radar plots showing percent total projections from each layer across HGEA-defined columns**. (A)** SUB_1, **(B)** SUB_2, **(C)** SUB_3, and **(D)** SUB_4. Axes represent target brain regions; values indicate percentage of total projections for each columnar and laminar group (Table 2).

**Table 1.** MPA single neuron reconstructions classified as HGEA cell types. Number of SUB neurons in each columnar (SUBdd-SUBdv) and laminar (Layer 1-4) group.

| HGEA Region | HGEA Layer |  |  |  | Total # |
| --- | --- | --- | --- | --- | --- |
| SUBdd | Layer 1 |  |  | Layer 4 | 167 |
|  | 137 |  |  | 30 |  |
| ProSUB |  |  | Layer 3 | Layer 4 | 111 |
|  |  |  | 59 | 52 |  |
| SUBv |  | Layer 2 | Layer 3 | Layer 4 | 273 |
|  |  | 40 | 138 | 95 |  |
| SUBvv |  | Layer 2 | Layer 3 | Layer 4 | 55 |
|  |  | 33 | 12 | 10 |  |
| SUBdv | Layer 1 |  |  | Layer 4 | 83 |
|  | 64 |  |  | 19 |  |
| Total # | 201 | 73 | 209 | 206 | 689 |

**Table 2.**
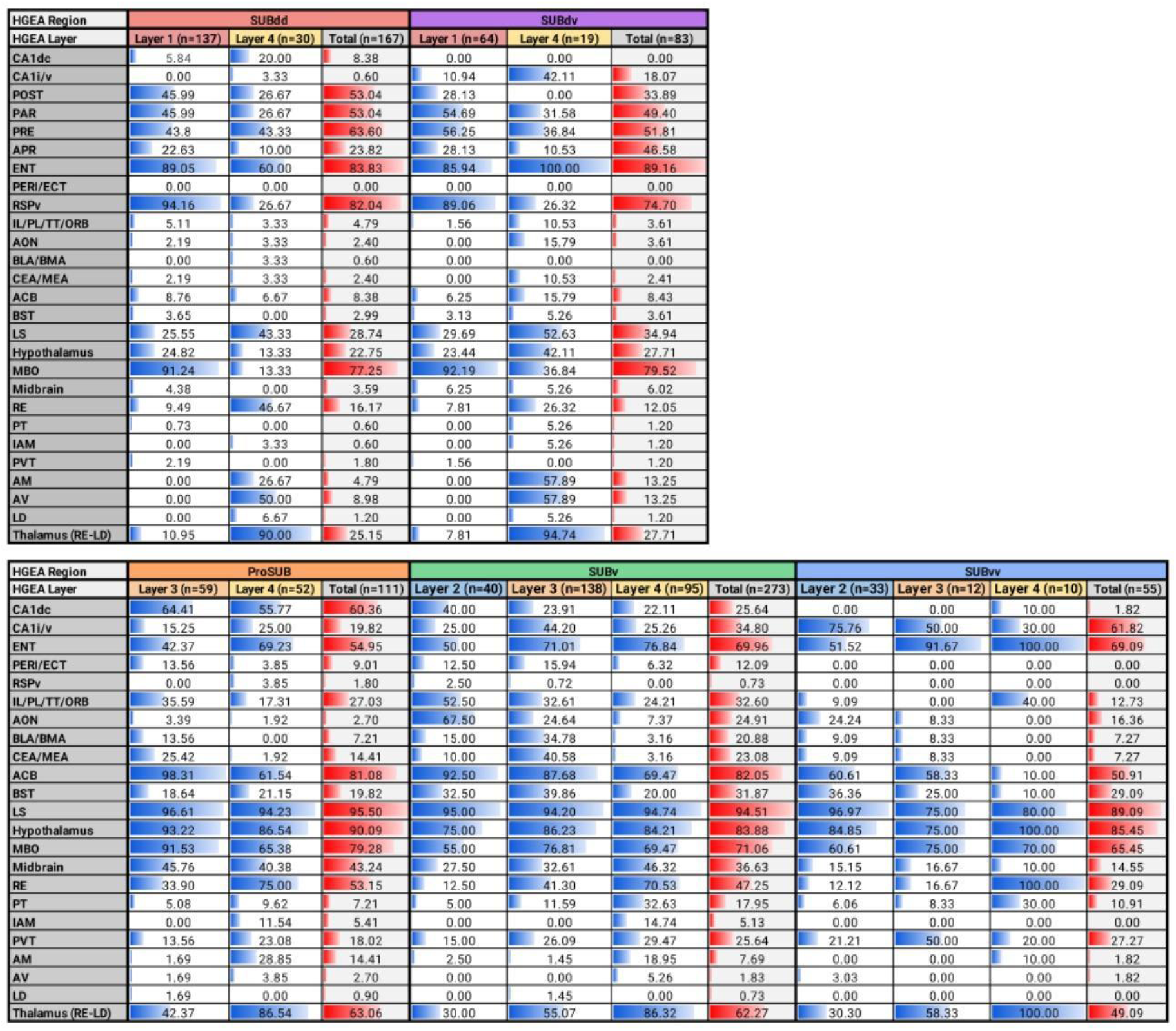
Percent of total neuron projections. Percent of total projections of each cell-type specific HGEA region (Total) and layers to various brain structures from top to bottom: CA1 dorsal of hippocampus, CA1 intermediate/ventral of hippocampus, entorhinal cortex, perirhinal/ectorhinal cortex, infralimbic/paralimbic/taenia tecta/orbital cortex, anterior olfactory nucleus, basal lateral and basal medial nuclei of the amygdala, central and medial nuclei of the amygdala, nucleus accumbens, bed stria terminalis, lateral septum, hypothalamus, mammillary body, midbrain, reuniens thalamic nucleus, paratenial thalamic nucleus, interanteromedial thalamic nucleus, paraventricular thalamic nucleus, anteromedial thalamic nucleus, anteroventral thalamic nucleus, anterodorsal thalamic nucleus, and lateral dorsal thalamic nucleus. Blue and red data bars are included for visual comparison between laminar and columnar groups.

These data bridge single-cell heterogeneity with population-level organizational patterns, revealing both common SUB connectivity profiles and rare cell-type specific variations within each HGEA group. Notably, these data also highlight SUB projections extending beyond classical hypothalamic targets to include substantial innervation of midbrain periaqueductal gray (PAG) and superior colliculus (SC) regions. While these descending pathways were first described by Köhler (1990), they have received limited attention in subsequent connectivity studies, and their relationship to HGEA laminar organization has not been previously characterized. In the following sections, we describe the connectivity features of each HGEA region and cell-type layer in detail.

### b. Dorsal Subiculum (SUBdd/dv)

Gene expression and connectivity analyses from the HGEA revealed two major cell-type layers within both SUBdd (n=167) and SUBdv (n=83): Layer 1, comprising primarily IT and ET neurons, and Layer 4, comprising primarily IT and CT neurons (**Fig. 6A**). Together, SUBdd and SUBdv constitute the bilaminar distal SUB, which extends along a curved dorsoventral trajectory from its rostral-dorsal position, corresponding to the classical dorsal SUB (SUBdd) to a caudal-ventral extension (SUBdv) where Layer 4 forms characteristic bulb-shaped enlargements. In the Allen CCFv3, this region corresponds broadly to the CCFv3 SUB subregion, though as discussed above, CCFv3 SUB boundaries do not appear to accurately align to the SUBdd/SUBdv boundaries as expected. Despite their shared bilaminar architecture, SUBdd and SUBdv exhibit complementary topographic projection patterns to shared target regions and are distinguished by their position along the hippocampal axis and differential inputs from dorsal vs. ventral CA1 regions.

Across all SUBdd and SUBdv neurons, the most common projections are to the entorhinal cortex (ENT; SUBdd = 89.1%, and SUBdv = 85.9%) (**Fig. 5G**) and mammillary body (MBO; SUBdd = 77.3% and SUBdv = 79.5%) (**Fig. 5C**). However, the most defining output signatures of the SUBdd and SUBdv are projections to the ventral retrosplenial cortex (RSPv), the subicular complex (POST, PRE, PAR), and anterior thalamic nuclei. These projections are not only highly represented but also distributed differently across layers rather than uniformly shared, as detailed in subsections below. Notably, septo-hypothalamic projections, which comparatively dominate the output profiles of ProSUB, SUBv, and SUBvv, are sparse in the SUBdd/dv, indicating a key connectivity distinction between the two major columnar architectures.

**Figure 5.**
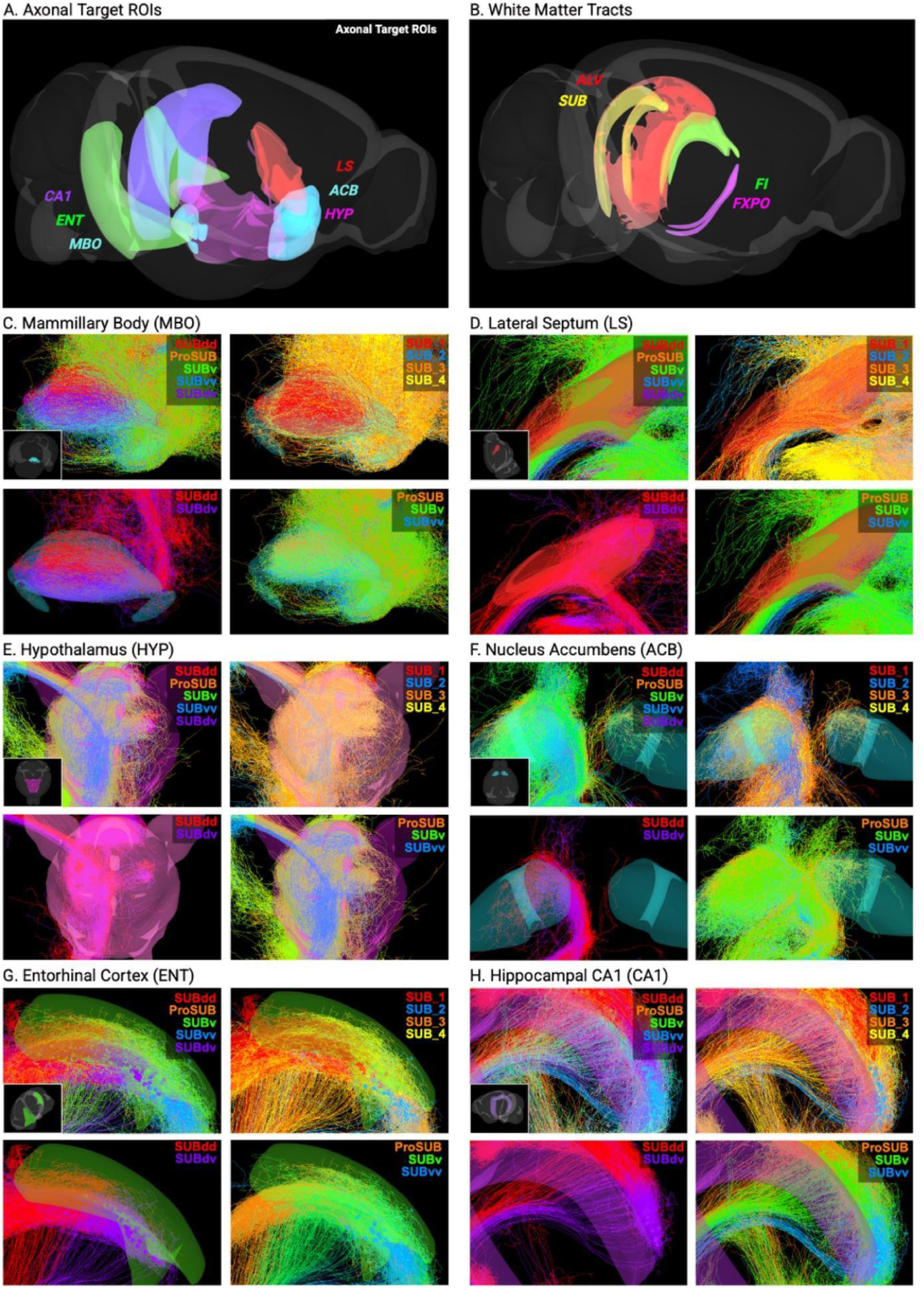
Brain regions that receive SUB axonal projections from all laminar and regional cell populations. **(A)** Overview of brain regions receiving projections from all SUB laminar (SUB_1–SUB_4) and regional (SUBdd, ProSUB, SUBv, SUBvv, SUBdv) populations. Highlighted regions include the mammillary body (MBO), lateral septum (LS), hypothalamus (HYP), nucleus accumbens (ACB), entorhinal cortex (ENT), and hippocampal CA1 (CA1). **(B)** SUB region with white matter tracts, including alveus (ALV), fimbria (FI), and postcommissural fornix (FXPO). **(C–H)** Representative projection patterns of HGEA laminar and regional SUB populations to individual target regions: mammillary body **(C)**, lateral septum **(D)**, hypothalamus **(E)**, nucleus accumbens **(F)**, entorhinal cortex **(G)**, and CA1 **(H)**. Colors correspond to SUB layers or regions as indicated. Insets show whole-brain reference of ROIs for spatial orientation.

#### SUBdd/SUBdv Layer 1 Cell Types

Layer 1 neurons are predominantly ET class neurons (SUBdd_1 = 94.2%, SUBdv_1 = 89.1%), with collateralized output to ipsilateral RSPv (**Fig. 6B**) and bilateral mammillary body (MBO; **Fig. 5C**). In both SUBdd_1 (n=137) and SUBdv_1 (n=64), more than 80% of Layer 1 neurons had collateralized projections to RSPv and MBO, exceeding prior estimates from dual retrograde tracing studies (Kinnavane et al., 2018). Notably, individual Layer 1 neurons send multiple separate axon collaterals to innervate various combinations of RSPv layers 2/3, 5, and/or 6 (although layer 2/3 is the primary target), with some neurons potentially sending two different collaterals to different portions of the same layer. The bifurcation point of these projections was highly variable: some axons traveled as a single fiber before splitting within RSPv, while others bifurcated near the soma and followed separate pathways to reach the same target.

**Figure 6.**
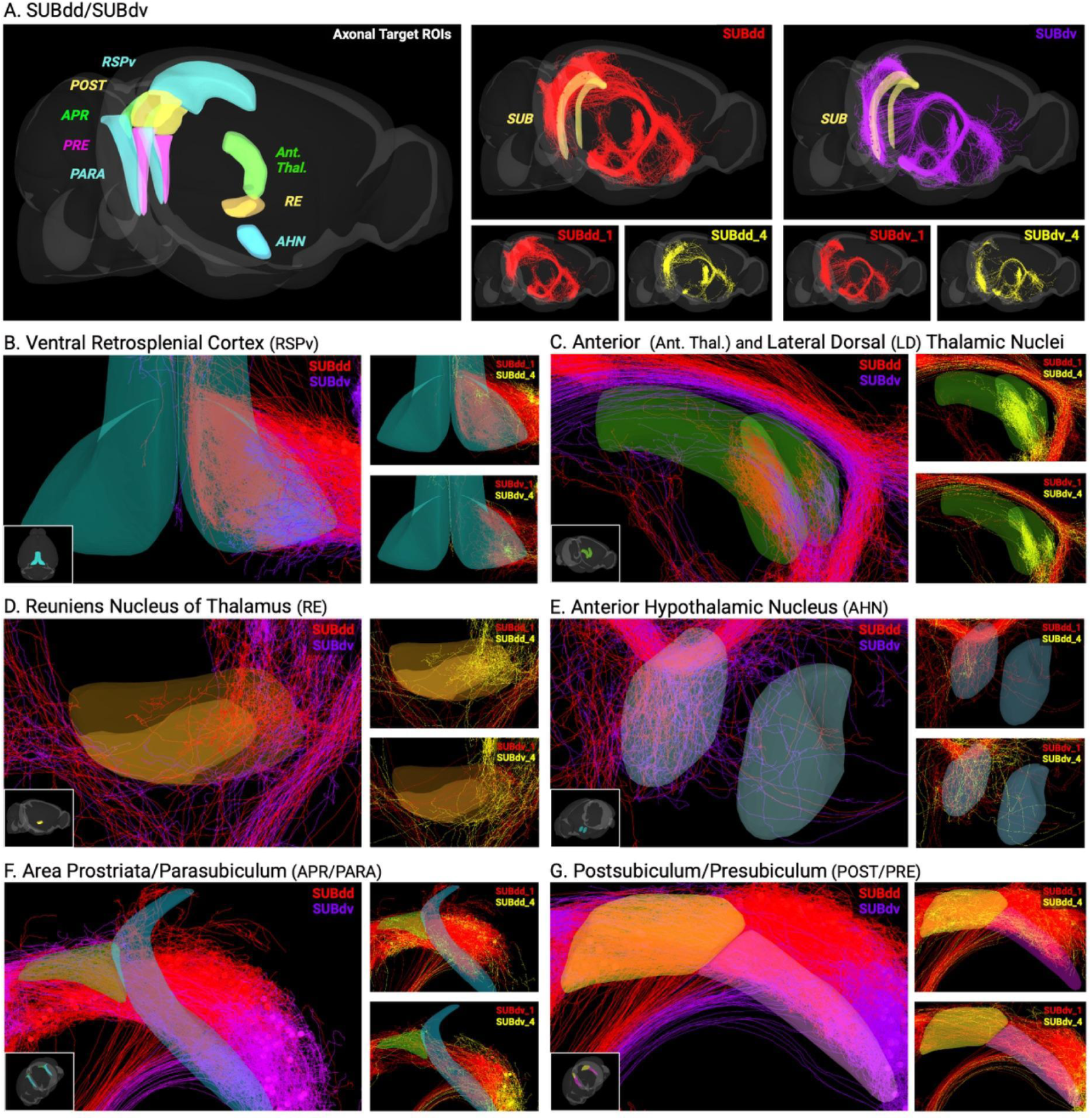
Differential projection patterns of dorsal subicular subdivisions (SUBdd and SUBdv). **(A)** Overview of brain regions receiving projections from SUBdd and SUBdv neuron populations. Colored regions indicate major target areas including ventral retrosplenial cortex (RSPv), anterior thalamic nuclei (Ant. Thal.) and laterodorsal thalamic nucleus (LD), reuniens nucleus (RE), anterior hypothalamic nucleus (AHN), area prostriata (APR), parasubiculum (PARA), postsubiculum (POST), and presubiculum (PRE). Representative whole-brain projection patterns for SUBdd (red) and SUBdv (magenta) are shown, with layer-specific examples below. **(B–G)** Region-specific projection patterns of SUBdd and SUBdv populations to individual target areas: **(B)** ventral retrosplenial cortex (RSPv), **(C)** anterior group of dorsal thalamic nuclei, including anteroventral (AV), anterodorsal (AD), anteromedial (AM), interanteromedial (IAM) thalamic nuclei and LD, **(D)** reuniens nucleus of the thalamus (RE), **(E)** anterior hypothalamic nucleus (AHN), **(F)** area prostriata/parasubiculum (APR/PARA), and **(G)** postsubiculum/presubiculum (POST/PRE). Insets show whole brain references of ROIs for spatial orientation. Colors correspond to subregional or laminar identities as indicated.

Layer 1 neurons also send collaterals to the postsubiculum (POST; SUBdd_1 = 46.0%, SUBdv_1 = 28.1%), presubiculum (PRE; SUBdd_1 = 44.5%, SUBdv_1 = 56.3%) (**Fig. 6G**), parasubiculum (PAR; SUBdd_1 = 46.0%, SUBdv_1 = 54.7%) (**Fig. 6F**), and entorhinal cortex (ENT; SUBdd_1 = 89.1%, SUBdv_1 = 85.9%) (**Fig. 5G**). ENT axon collaterals were relatively short, innervating layers 5 and 6 of the caudal dorsomedial ENT (Van Groen, 2001), a region containing grid cells with the highest spatial grid scale (Brun et al., 2008). Deep-layer ENT neurons have in turn been shown to innervate superficial ENT layers (Simonsen et al., 2022), forming a feedback loop that may influence ENT-hippocampal input connectivity. Notably, only 22.8% of SUBdd_1 and 20.3% of SUBdv_1 neurons lacked output to all three of POST, PRE, and PAR, indicating that collateralization within the subicular complex is the rule rather than the exception. In SUBdd_1, collateral combinations were roughly evenly distributed across categories (9–15% each). In SUBdv_1, projections were biased toward PRE and PAR over POST, suggesting that SUBdv_1 output is preferentially directed toward PRE and PAR circuits.

The area prostriata (APR), a region implicated in processing fast motion in the peripheral visual field (Mikellidou et al., 2017), was recently identified in mice and shown to receive inputs from primary visual cortex and dorsal, but not ventral, SUB (Hu et al., 2020; Lu et al., 2020). We found that 22.6% of SUBdd_1 and 26.6% of SUBdv_1 neurons projected to APR (**Fig. 6F**), each with a clearly branched axon collateral. Because the APR in turn projects back to POST and PRE (Chen et al., 2021), these Layer 1 neurons appear to participate in a reciprocal visuospatial circuit linking the subicular complex and APR. A small proportion of Layer 1 neurons also sent back projections to other hippocampal structures (**Fig. 5H**), most commonly CA1, with one neuron targeting the dentate gyrus (DG), typically as a short branch off the axon traveling through the alveus (SUBdd_1 = 5.8%, SUBdv_1 = 10.9%).

#### SUBdd/SUBdv Layer 4 Cell Types

SUBdd and SUBdv Layer 4 neurons (SUBdd_4 n=30, SUBdv_4 n=19) are primarily classified as CT-type neurons targeting the reuniens nucleus (RE; **Fig. 6D**), anteroventral nucleus (AV), anterodorsal nucleus (AD), anteromedial nucleus (AM), and laterodorsal (LD) thalamic nuclei (**Fig. 6C**). However, many of these neurons can have collateral projections to a variety of extra-telencephalic brain regions. Retrograde tracer injections into these thalamic regions revealed that the Layer 4 of the SUBdd and SUBdv contains a specialized sublayer primarily containing the AV-, AD-, and LD-projecting CT subpopulations where Layer 4 becomes thicker at its distal ends (Bienkowski et al., 2018). In contrast, the RE-projecting subpopulations are located in the deeper parts of Layer 4. To simplify the naming of these sublayers, we will refer to the deeper RE-projecting sublayer as Layer 4.2 and the superficial sublayer containing other cell types as Layer 4.1. At the most caudal level of the SUB (HGEA 96) where only Layer 4 exists, the SUBdd and SUBdv Layer 4.1 sublayers become continuous along the border with the presubiculum (PRE).

Within the MPA dataset, RE-projecting neurons (Layer 4.2) comprised 46.7% of SUBdd_4 and 26.3% of SUBdv_4 neurons, confirming this population as largely distinct from other Layer 4 subtypes. Some RE-projecting neurons solely targeted the RE whereas others had collateral projections to septo-hypothalamic regions (**Fig. 6E**). In SUBdd_4, AV-projecting neurons were the most prevalent subtype (50.0%), while SUBdv_4 showed a broader thalamic targeting profile with substantial proportions of both AV-projecting (57.9%) and AM-projecting (57.9%) neurons alongside the RE-projecting population. Beyond their thalamic targets, SUBdv_4 neurons showed more extensive collateralization than SUBdd_4, with 26.3% projecting to RSPv (**Fig. 6B**) and additional neurons targeting septo-hypothalamic regions.

Notably, SUBdv_4 projections to RSPv innervate deep cortical layers, in contrast to SUBdd_1 and SUBdv_1 neurons which target RSPv layer 2.

### c. Prosubiculum (ProSUB)

The ProSUB (n=111) is a dorsal region that contains two cell-type layers, Layer 3 (n=59), comprising primarily ET neurons, and Layer 4 (n=52), comprising primarily IT and CT neurons, distinguishing it from the SUBv/SUBvv due to its lack of superficial layer 2 (**Fig. 7A**).

**Figure 7.**
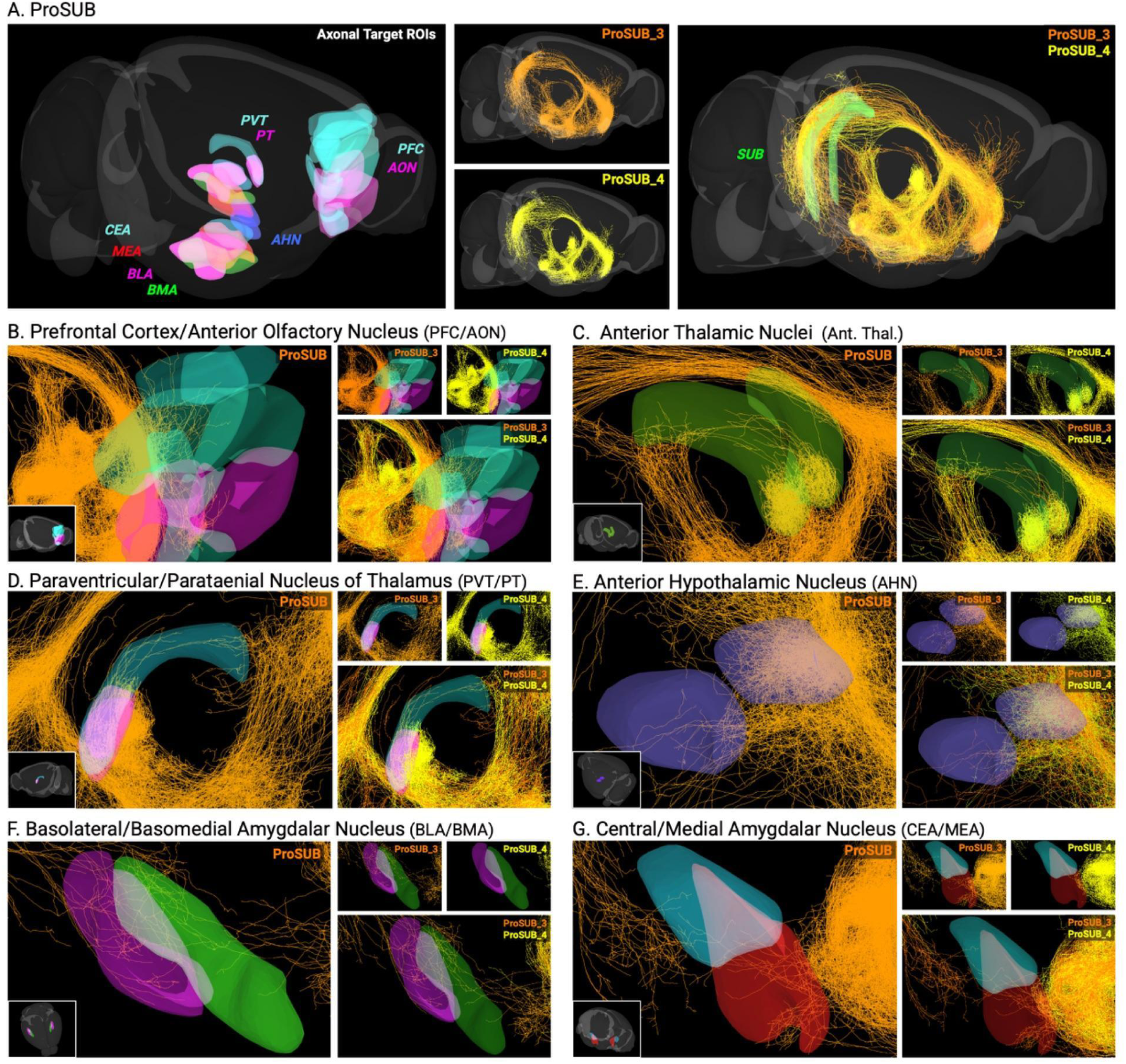
Projection targets of ProSUB laminar populations. **(A)** Overview of brain regions receiving projections from ProSUB neuron populations. Colored regions indicate major forebrain targets including prefrontal cortex/anterior olfactory nucleus (PFC/AON), anterior thalamic nuclei and laterodorsal thalamic nucleus (LD), paraventricular/parataenial thalamic nuclei (PVT/PT), anterior hypothalamic nucleus (AHN), basolateral/basomedial amygdala (BLA/BMA), and central/medial amygdala (CEA/MEA). Representative whole-brain projection patterns for ProSUB_3 (orange) and ProSUB_4 (yellow) are shown at right. **(B–G)** Region-specific projection patterns of ProSUB laminar populations to individual target regions: **(B)** PFC/AON, **(C)** anterior group of dorsal thalamic nuclei, **(D)** PVT/PT, **(E)** AHN, **(F)** BLA/BMA, and **(G)** CEA/MEA. Insets show whole brain references of ROIs for spatial orientation. Colors correspond to laminar identity as indicated.

In the HGEA, the ProSUB first appears rostrally as the CA1/SUB border shifts lateral to the perpendicular of the dentate gyrus blade tip, positioning it proximal to SUBdd between CA1 and the distal SUB. At its caudal end, the ProSUB terminates at HGEA level 91, where layer 2 rises dorsally to meet the SUBdd border and the region transitions into SUBv. In the Allen CCFv3, the prosubiculum is defined more broadly, encompassing nearly the entire ventral SUB area that the HGEA defines as SUBv and parts of the SUBvv, with no separate ventral area. While the ProSUB and SUBv share some similar output targets and their layers 3 and 4 are anatomically continuous, we distinguish them as separate structures based on their fundamentally different laminar architectures, bilaminar (layers 3 and 4) versus trilaminar (layers 2, 3, and 4), and the distinct connectivity profiles described below.

Across all ProSUB layers, the most prevalent shared projections are to the lateral septum (LS; 95.5%), hypothalamus (90.1%), ACB (81.1%), and MBO (79.3%), with substantial midbrain projections (43.2%) (**Fig. 5C-F**). In striking contrast to SUBdd/dv, RSPv connectivity is nearly absent (1.8%), representing a fundamental shift in the cortical output profile. ENT projections are well-represented overall (55.0%) but are unevenly distributed across layers (ProSUB_3 = 42.4%, ProSUB_4 = 69.2%; **Fig. 5G**). Notably, ProSUB targets deep layers of the lateral ENT, which is topographically distinct from the caudal dorsomedial ENT region favored by SUBdd/dv, and 9% of ProSUB neurons extend collaterals further to the perirhinal/ectorhinal cortex (PERI/ECT). Additional projections include the prefrontal cortex (IL/PL/TT/ORB; 27.0%; **Fig. 7B**), bed nucleus of the stria terminalis (BST; 19.8%), basolateral/basomedial amygdala (BLA/BMA; 7.2%; **Fig. 7F**), and central/medial amygdala (CEA/MEA; 14.4%; **Fig. 7G**).

Importantly, non-canonical back projections to the hippocampus are far more prevalent in ProSUB than in SUBdd/dv, with 60.4% of ProSUB neurons sending collaterals to dorsal CA1 and an additional 19.8% extending collaterals to the more distant CA1i/CA1v regions (**Fig. 5H**).

#### ProSUB Layer 3 Cell Types

ProSUB Layer 3 neurons are primarily ET-class neurons and are most clearly distinguished from Layer 4 by more prevalent projections to the prefrontal cortex (35.6% vs 17.2%; **Fig. 7B**), amygdala (BLA/BMA = 13.5% vs. 0%; CEA/MEA=25.4% vs. 1.9%; **Fig. 7F,G**), and PERI/ECT (13.6% vs. 3.9%). Although ACB and MBO projections are common across both layers, they are approximately 30% more prevalent in Layer 3, consistent with the ET-class designation (ACB = 98.3%, MBO = 65.4%).

#### ProSUB Layer 4 Cell Types

ProSUB Layer 4 neurons are primarily CT-class neurons and target the reuniens nucleus (RE) at the highest rate of any SUB Layer 4 population (75.0%), with many of these neurons also sending collateral projections to a variety of telencephalic and extra-telencephalic regions. Unlike SUBdd/dv Layer 4, ProSUB_4 shows a marked shift away from SUBdd/dv_4-like anterior thalamic targeting (**Fig. 7C**), with very low rates of AV- (3.9%), AD-(0.0%), and LD-projecting (0.0%) neurons. Instead, interanteromedial (IAM; 11.5%) and anteromedial (AM; 28.9%) thalamic projections are observed. Projections to midline thalamic structures like paraventricular thalamic nucleus (PVT; 23.1%) and paratenial nucleus (PT; 9.6%) (**Fig. 7D**) projections are also substantially higher than in SUBdd_4 (PVT = 0%, PT = 0%) or SUBdv_4 (PVT = 0%, PT = 5.3%). This thalamic targeting profile positions ProSUB_4 as an intermediate CT population, sharing the RE-dominant motif common to all SUB Layer 4 subtypes while transitioning toward the midline and paraventricular thalamic targets that characterize SUBv/SUBvv Layer 4.

### d. Ventral Subiculum (SUBv/SUBvv)

The SUBv (n=273) and SUBvv (n=55) are both trilaminar structures containing Layers 2, 3, and 4, distinguishing them from the bilaminar ProSUB and SUBdd/dv (**Fig. 8A**). SUBv occupies the intermediate ventral SUB, while SUBvv represents its most ventral extent, where the relative size and shape of the layers shift as they approach the ventral tip. In the Allen CCFv3, part of this ventral SUB area has been re-designated as the hippocampal amygdala transition area (HATA). However, our recent work demonstrates that this area reflects the anatomical transition of separate CA1 and SUB laminar cell types rather than a discrete structural region (Pachicano et al., 2025). Because MPA neurons registered to the CCFv3 showed profiles consistent with SUBv/SUBvv, we assigned HATA neurons to their corresponding SUB layers and treated them as part of the SUBv/SUBvv population throughout this analysis.

**Figure 8.**
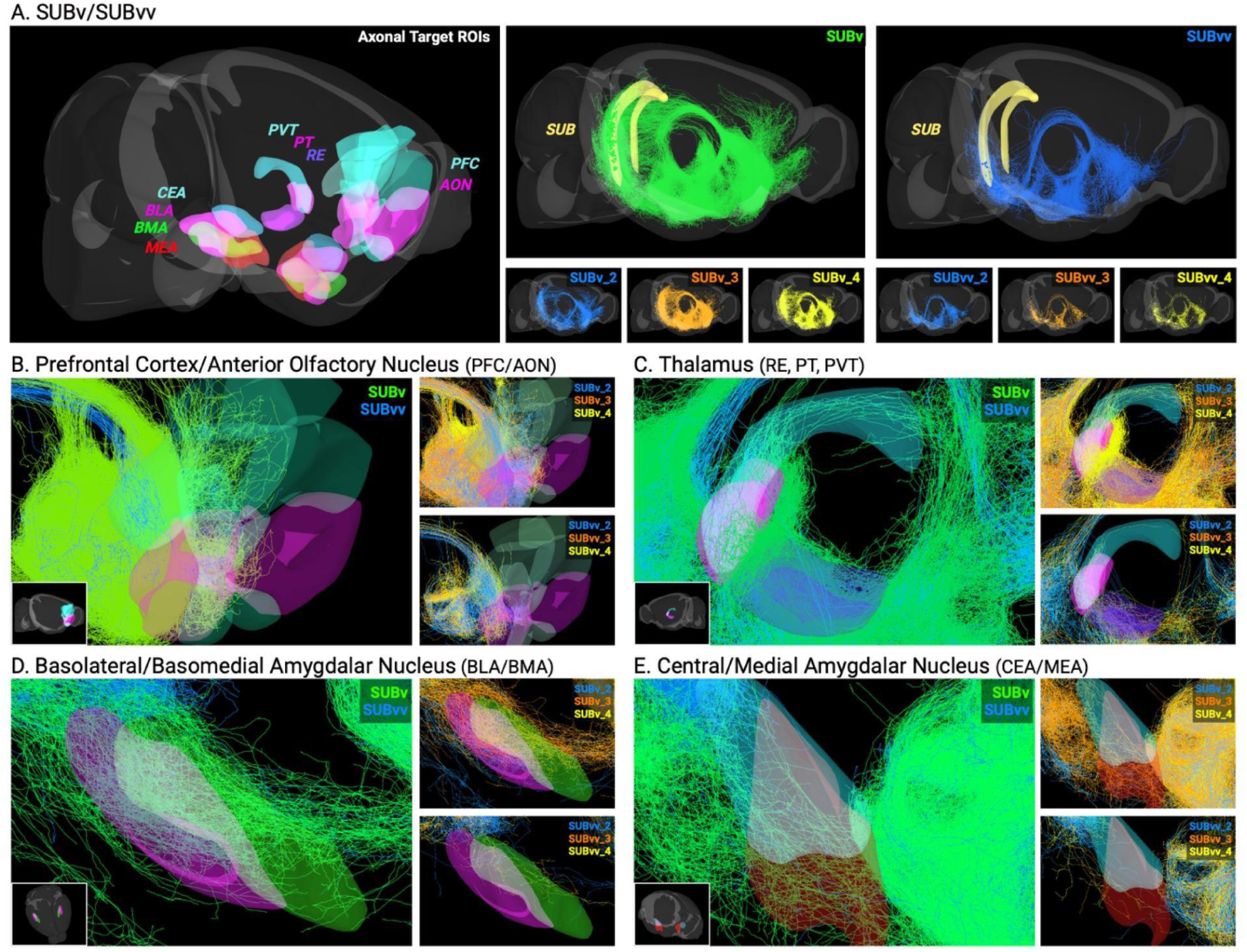
Projection targets of ventral subicular subdivisions (SUBv and SUBvv). **(A)** Overview of brain regions receiving projections from ventral subicular populations (SUBv and SUBvv). Colored regions indicate major forebrain targets including prefrontal cortex/anterior olfactory nucleus (PFC/AON), thalamic nuclei (reuniens, paraventricular, and parataenial; RE/PT/PVT), basolateral and basomedial amygdala (BLA/BMA), and central and medial amygdala (CEA/MEA). Representative whole-brain projection patterns for SUBv (green) and SUBvv (blue) are shown, with layer-specific examples displayed below. **(B–E)** Region-specific projection patterns of SUBv and SUBvv populations to individual target regions: **(B)** PFC/AON, **(C)** thalamus (RE/PT/PVT), **(D)** BLA/BMA, and **(E)** CEA/MEA. Insets show whole brain references of ROIs for spatial orientation. Colors correspond to subregional or laminar identities as indicated.

Across both SUBv and SUBvv, the most prevalent shared projections are to the lateral septum (LS; SUBv = 94.5%, SUBvv = 89.1%), hypothalamus (SUBv = 83.9%, SUBvv = 85.5%), ACB (SUBv = 82.1%, SUBvv = 50.9%), and MBO (SUBv = 71.1%, SUBvv = 65.5%; **Fig. 5C–F**).

ENT projections are also well-represented across both regions (SUBv = 70.0%, SUBvv = 69.1%; **Fig. 5G**). However, they exhibit a laminar bias as ACB-projecting neurons are more prevalent in superficial layers (2 and 3) while ENT-projecting neurons predominate deeper layers (3 and 4). In contrast to SUBdd/dv, RSPv projections are nearly absent across both regions, while the high prevalence of septo-hypothalamic and limbic outputs marks SUBv/SUBvv as the primary SUB conduit to motivational and neuroendocrine networks.

#### SUBv/SUBvv Layer 2 Cell Types

SUBv/SUBvv Layer 2 neurons are primarily IT-class neurons and are uniquely distinguished from other SUB layers by two defining output features identified in the HGEA: strong projections to the anterior olfactory nucleus (AON) and local collateralization along the CA1-SUB hippocampal axis. In the HGEA, Layer 2 neurons were characterized as a specialized population transferring information along the transverse axis from SUBvv toward SUBv, complementary to the reverse directionality seen in Layer 4 (SUBv → SUBvv). In addition, both of these layers have strong back projections to CA1, with SUB_2 having projections to CA1v and SUB_4 having projections CA1vv. In the MPA dataset, back projections to CA1 were prevalent in both SUBv_2 (CA1dc = 40.0%, CA1i/v = 25.0%) and SUBvv_2 (CA1i/v = 76.0%), confirming non-canonical hippocampal interconnectivity that is specific to this layer (Xu et al., 2016), **Fig. 5H**).

AON projections were strongly Layer 2-enriched, with 67.5% of SUBv_2 and 24.2% of SUBvv_2 neurons projecting to AON, compared to substantially lower rates in Layer 3 (SUBv = 24.6%, SUBvv = 8.3%) and Layer 4 (SUBv = 7.4%, SUBvv = 0%; **Fig. 8B**). The lower AON projection rate in SUBvv_2 may partly reflect sparse sampling, as AON-projecting neurons appear clustered at the ventral tip of Layer 2. Notably, 52.5% of SUBv_2 neurons had collaterals that extended dorsally from AON into the prefrontal cortex (infralimbic (IL), prelimbic (PL) taenia tecta (TT), and orbitofrontal (ORB) cortex), establishing a direct SUB-to-prefrontal pathway that is unique to this layer.

#### SUBv/SUBvv Layer 3 Cell Types

SUBv/SUBvv Layer 3 neurons are primarily ET-class neurons and are distinguished from other SUB layers by more prominent projections to amygdalar nuclei and midline thalamic nuclei. Within the amygdala, 34.8% and 40.6% of SUBv_3 neurons project to the BLA/BMA and CEA/MEA respectively (**Fig. 8D,E**), while SUBvv_3 has substantially fewer amygdala projections (8.33% to both BLA/BMA and CEA/MEA). Many of these amygdala-projecting axons can take one or both of two routes: a direct anterior projection route through the ventral SUB, or as a longer collateral branching from the fornix, traveling laterally through the hypothalamus.

Beyond the amygdala, SUBv_3 neurons also strongly target midline thalamic nuclei which is atypical of Layer 3 cell types elsewhere in the SUB (**Fig. 8C**). While 41.3% of SUBv_3 neurons project to the RE, this decreases substantially to 16.7% in SUBvv_3. In contrast, both SUBv_3 (26.1%) and SUBvv_3 (50.0%) show strong projections to the PVT, indicating a pronounced bias towards SUBvv_3 for this pathway. Although these thalamic projections in Layer 3 are notable, they remain less prevalent than the thalamic projections of SUBv/SUBvv Layer 4.

#### SUBv/SUBvv Layer 4 Cell Types

SUBv/SUBvv Layer 4 neurons are primarily CT-class neurons and also show the highest degree of collateralized output to the ENT of any SUB layer (**Fig. 5G**). Like other Layer 4 SUB neurons, the RE is the most prevalent thalamic target, innervated by 70.5% of SUBv_4 and 100% of SUBvv_4 neurons. SUBv/SUBvv Layer 4 neurons also strongly collateralize to the PVT (SUBv_4 = 29.5%; SUBvv_4 = 20.0%) and PT (SUBv_4 = 32.6%; SUBvv_4 = 30.0%), with axons typically entering the PVT at its rostral end and continuing caudally at varying lengths (**Fig. 8C)**. Notably, a subset of neurons send an axon into the PVT from the caudal end as a collateral of the ascending branch targeting the midbrain PAG/SC. Via these routes, 46.3% of SUBv_4 and 10.0% of SUBvv_4 neurons projected to the midbrain (**Fig. 9**), a finding discussed in the midbrain section below.

**Figure 9.**
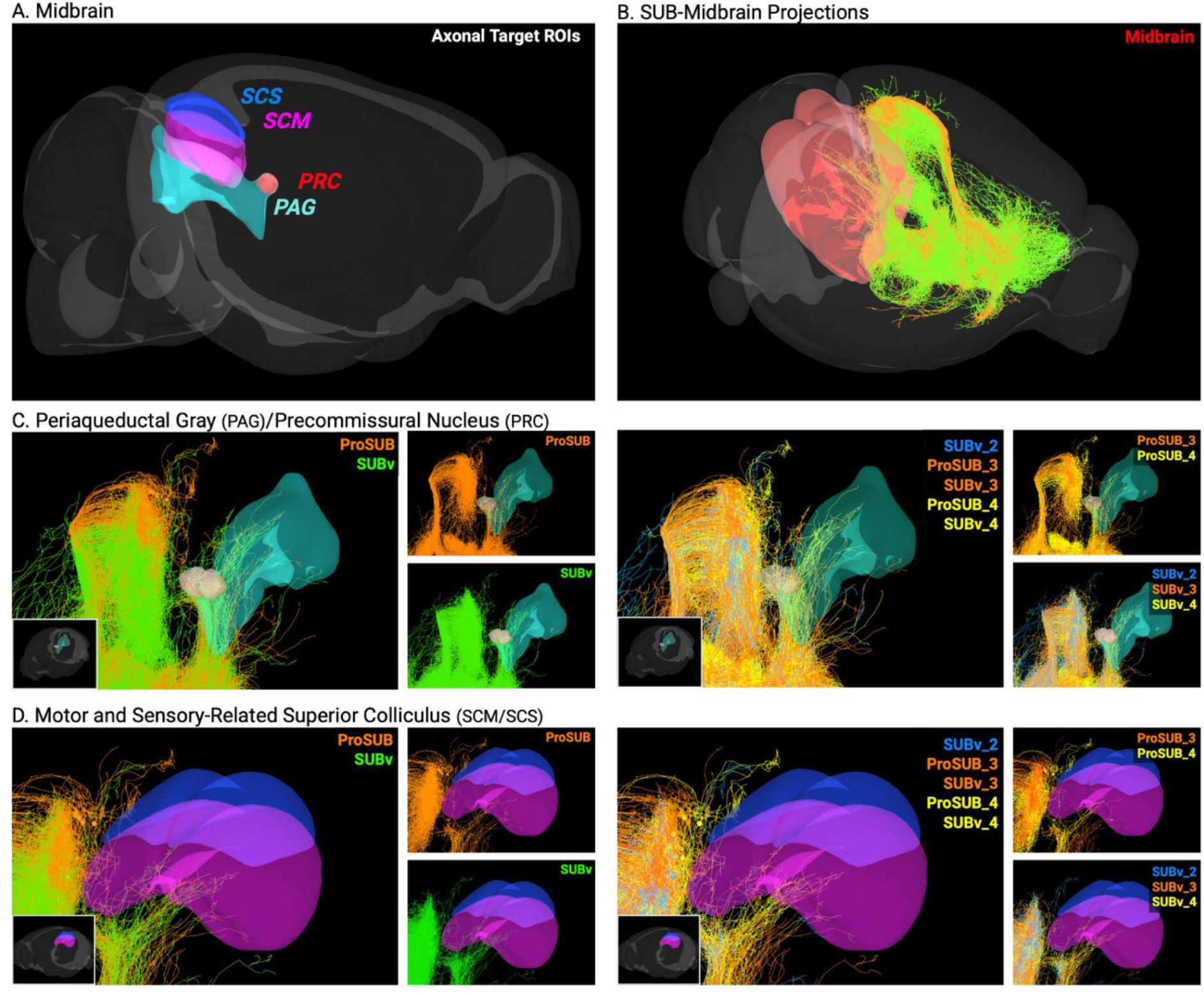
Columnar and laminar projections to midbrain structures. **(A)** Overview of midbrain brain regions receiving projections from ProSUB and SUBv neuron populations. Colored regions indicate major midbrain regions, including motor and sensory- related superior colliculus (SCM/SCS), precommissural nucleus (PRC), and the periaqueductal gray (PAG). **(B)** SUB to midbrain projections. **(C–D)** Region-specific projection patterns of ProSUB and SUBv populations to individual target regions: **(C)** PAG/PRC and **(D)** SCM/SCS. Insets show whole brain references of ROIs for spatial orientation. Colors correspond to subregional or laminar identities as indicated.

Across all five HGEA-defined SUB regions, the MPA single-neuron reconstructions reveal both the organizing principles that unify each cell-type group and the heterogeneity that distinguishes individual neurons within them. In the following sections, we discuss how these findings confirm, extend, and in some cases substantially revise our understanding of SUB cell type connectivity and its contributions to brain-wide networks.

## IV: Discussion

### a. Laminar and columnar organization of the SUB at single-cell resolution

Through virtual tract tracing of 689 MPA single-neuron reconstructions, this study provides a comprehensive single-cell validation of the laminar-columnar framework of the SUB established by the HGEA. Individual SUB neurons exhibit projection patterns that closely match population-level predictions based on retrograde tracing and gene expression studies (Bienkowski et al., 2018, 2021). When visualized in 3D space, connectivity-based classification alone recapitulates the spatial organization of HGEA defined layers and columns, demonstrating that the gene expression boundaries correspond to functionally distinct organizational units at the single-cell level. While the large majority of neurons conform to layer-specific projection motifs, single-cell data also reveals variation within each cell-type group that extends beyond what population-level tracing could resolve, providing new insights into the heterogeneity and collateralization principles that distribute SUB output throughout the brain.

Underlying this classification are potential reconstruction errors, registration issues that arise from registering projectome data from multiple brains into the 3D CCFv3 atlas space, and the imperfect alignment between HGEA-defined cell-type boundaries and CCFv3 anatomical delineations. While CCFv3 SUB corresponds primarily to HGEA SUBdd/SUBdv and CCFv3 ProS to HGEA ProSUB, we identified neurons with projection patterns characteristic of all four HGEA layers within CCFv3 SUB boundaries. This likely reflects both genuine differences in how the two frameworks define regional boundaries and spatial imprecision inherent in registering individual MPA brains to the CCFv3 reference atlas. Notably, MPA neurons registered as CCFv3 ProS neurons likely also include all 4 layers and would include more neurons from the HGEA SUBvv that are undersampled here. Irrespective of the atlas boundaries and simply focusing on the 3D space, the classification of SUB MPA neurons as laminar cell types shows remarkable similarity to the 3D HGEA model (compare Fig. 2A vs Fig. 1A). However, the CCFv3 HATA region presents an additional challenge: while Ding et al. (2020) characterized HATA as a discrete structural region with unique transcriptomic identity, our recent work suggests it instead represents a transitional zone where CA1v/vv and SUBv/vv laminar cell types converge (Pachicano et al., 2025), and MPA neurons registered to HATA showed connectivity profiles consistent with SUBv/SUBvv, supporting their inclusion in that population here. Thus, misalignment between frameworks has practical implications for cross-study comparisons where data registered to CCFv3 anatomical boundaries may inadvertently sample from different combinations of HGEA layers.

Beyond these atlas-level organizational principles, single-cell data reveals structured variation within each HGEA group that reflects genuine cell-type heterogeneity rather than noise. Unlike transcriptomic studies where heterogeneity has been characterized primarily through gene expression, our analysis quantifies it through connectivity, which is an approach that directly reflects functional output diversity. Consistent with this, Campbell et al. (2026) recently demonstrated that SUB projection neurons exhibit variable dendritic and axonal complexity, such that morphological heterogeneity within the SUB is itself an organizational feature characteristic of projection-specific input-output relationships. Rather than representing random complexity, we believe this heterogeneity is a key organizational feature of brain circuits, enabling flexible and context-dependent network engagement rather than the rigid, homotypic activation that would result from populations of identically-wired neurons.

The SUBdd_1 population illustrates this principle clearly. Approximately 90% of SUBdd_1 neurons have an axon collateral that targets the subicular complex (POST, PRE, or PAR), APR, or ENT, which are brain regions strongly implicated in visuospatial navigation (Boccara et al., 2010; Hafting et al., 2005; Peyrache et al., 2017; S.-Y. Zhang et al., 2022).

However, the specific combination of targets innervated by any individual neuron is highly variable. Despite this variability, SUBdd_1 neurons provide roughly equivalent total input to POST, PRE, and PAR, and APR projections consistently appear as a collateral of subicular complex projections rather than as a sole target. This suggests that while individual neurons vary in precise combinations of regions they target, the population as a whole maintains structured and balanced output to this visuospatial circuit. Understanding these combinatorial principles for axon collateralization, or the rules that constrain which combinations of targets co-occur and which do not, is essential for linking single-cell connectivity to population-level dynamics.

### b. CT/IT/ET framework applied to SUB cell types

Previous studies established a tripartite classification of cortical projection neurons, including intratelencephalic (IT) neurons projecting to cortex and striatum, extratelencephalic (ET) neurons to subcortical regions, and corticothalamic (CT) neurons to the thalamus. Despite a great diversity of cortical neuron cell types, these general classifications have been proven to be applicable across neocortical areas (Shepherd, 2013; Harris et al., 2019; Winnubst et al., 2019; Yao et al., 2023). Although the SUB is regarded as an allocortical structure, SUB pyramidal cell type layers exhibit analogous projection classes and later transcriptomic studies substantiated this view (Yao et al., 2023). Our observations of the MPA single-neuron projectome data further validates this perspective, but also demonstrates that SUB neurons don’t follow the rules as strictly as neocortical neurons (Bienkowski et al., 2018; Gao et al., 2026). For example, the high percentage of SUB neurons with subcortical projections to hypothalamus or MBO would, by strict definition, classify nearly all as ET neurons. Indeed, thalamic-projecting CT neurons and cortical-projecting IT neurons without subcortical collaterals exist but are more rare when compared to the whole population. Still, given a more relaxed projection-based definition and relationships to neocortical transcriptomic profiles, Layer 6-like SUB_4 neurons align to the CT-class across all five SUB regions, consistently targeting RE and other thalamic nuclei whereas SUB_1, SUB_2, and SUB_3 neurons closely resemble mixed IT or ET classes. SUB_1 and SUB_2 neurons show the closest conformation to the IT-class with projections to cortical structures and striatal-like LS and ACB. Finally, SUB_3 neurons conform to the ET-class as they have increased subcortical outputs to the amygdala and midbrain.

Evolutionary studies of the brain suggest that the hippocampus and SUB were some of the earliest cortical areas to evolve in the brain (Woych et al., 2022), so their strong association with hypothalamus and other subcortical structures should be expected as there were few other cortical areas to connect with in the early vertebrate brain. This suggests that the subcortical connectivity was originally inherent to early cortical cell types and that ET and IT neurons as segregated classes likely diverged later in evolution as cortical regions expanded. Thus, the SUB may be an important site for understanding cortical brain architecture where the IT/ET/CT cell type framework may have first emerged and where laminar and columnar organizational principles converged to form the template for canonical cortical microcircuits (although further study of synaptic connections between SUB cell types is needed) (Bastos et al., 2012). Overall, a conserved organizational principle of cortical output extends beyond the neocortex to allocortical structures, even as the specific downstream circuits diverge considerably (**Fig. 10**).

**Figure 10.**
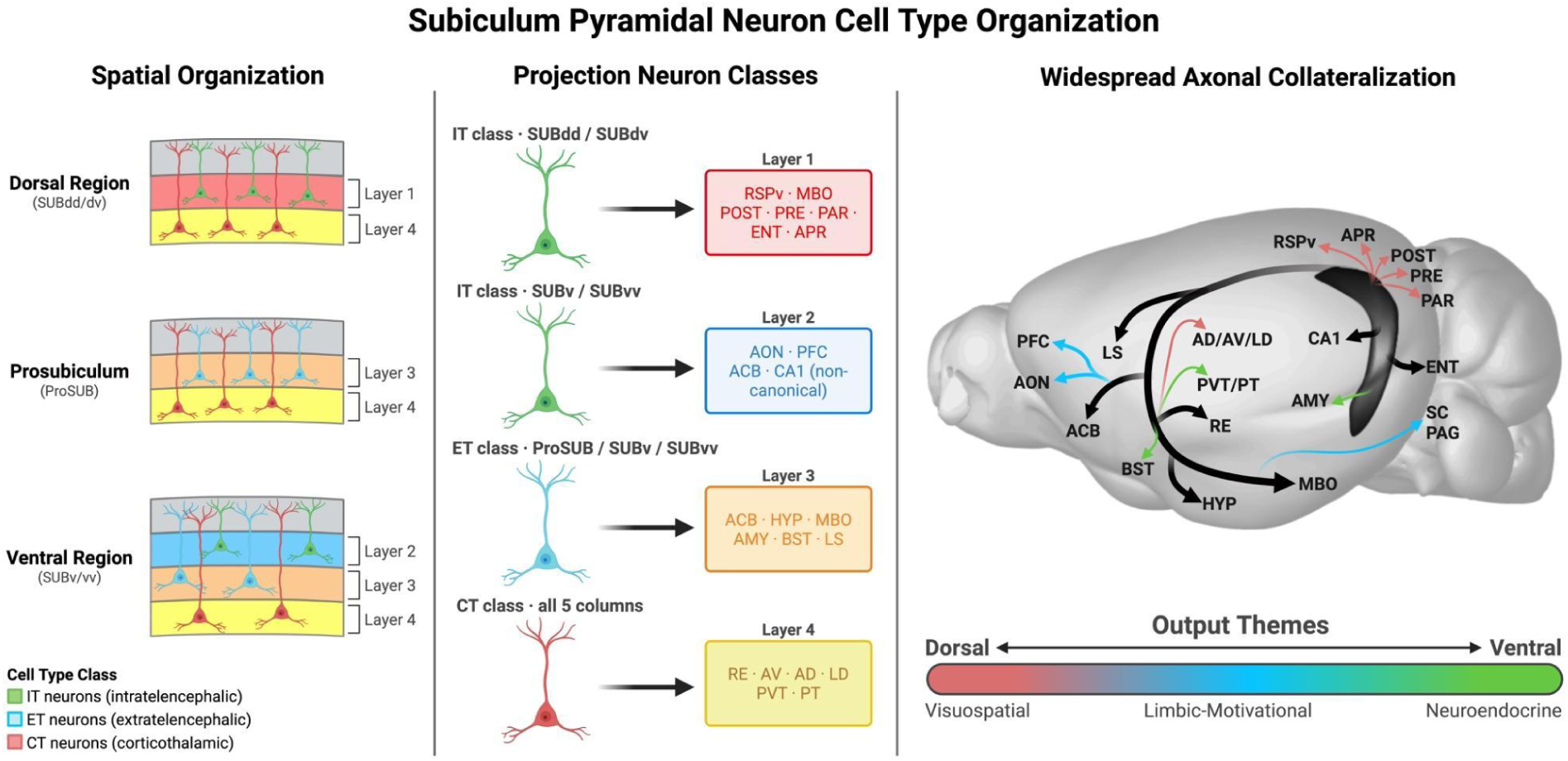
Subiculum Pyramidal Neuron Cell Type Organization. (Left) Spatial organization of the HGEA-defined SUB laminar cell types across the dorsal (SUBdd/dv), prosubiculum (ProSUB), and ventral (SUBv/vv) regions. Layer 1 and Layer 2 neurons are intratelencephalic (IT; green), Layer 3 neurons are extratelencephalic (ET; blue), and Layer 4 neurons are corticothalamic (CT; red). **(Middle)** Cell type-specific projection targets for each laminar group based on highest proportion (Table 2). Layer 1 IT neurons (SUBdd/dv) project to the ventral retrosplenial cortex (RSPv), mammillary body (MBO), postsubiculum (POST), presubiculum (PRE), parasubiculum (PAR), entorhinal cortex (ENT), and area prostriata (APR). Layer 2 IT neurons (SUBv/vv) project to the anterior olfactory nucleus (AON), prefrontal cortex (PFC), nucleus accumbens (ACB), and hippocampal CA1 region via non-canonical back projections. Layer 3 ET neurons (ProSUB/SUBv/SUBvv) project to the ACB, hypothalamus (HYP), MBO, amygdala (AMY), bed nucleus of the stria terminalis (BST), and lateral septum (LS). Layer 4 CT neurons (all five columns) project to the nucleus reuniens (RE), anteroventral thalamic nucleus (AV), anterodorsal thalamic nucleus (AD), laterodorsal thalamic nucleus (LD), paraventricular thalamic nucleus (PVT), and paratenial nucleus (PT). **(Right)** Whole-brain schematic illustrating the widespread axon collateralization of SUB cell types and their output projection themes along the dorsoventral axis, spanning visuospatial (dorsal), limbic-motivational (intermediate), and neuroendocrine (ventral) networks.

### c. Dorsal Subiculum (SUBdd/dv)

Prior population tracing and gene expression studies established that SUBdd and SUBdv Layer 1 neurons project to the subicular complex (POST, PRE, and PAR), RSPv, ENT, and MBO, while Layer 4 neurons predominantly target the anterior thalamic nuclei, LD, and RE (Bienkowski et al., 2018). Classical tracing demonstrated that these projections are topographically organized rather than uniform, with SUBdd and SUBdv showing complementary biases across shared targets (Swanson & Cowan, 1975), and dual retrograde tracing further suggested that approximately half of dorsal SUB neurons collateralize to both RSPv and MBO, implicating a shared circuit for spatial memory consolidation (Kinnavane et al., 2018; Umaba et al., 2021). The MPA single-neuron reconstructions confirm and substantially extend these findings. RSPv/MBO dual collateralization in Layer 1 neurons exceeds prior retrograde tracing estimates, with more than 80% of both SUBdd_1 and SUBdv_1 neurons projecting to both targets. This is consistent with recent fMOST-based single-neuron projectome data in the dorsal SUB (J. Zhang et al., 2025). Furthermore, SUBdd_1 was biased toward POST and caudal PAR, SUBdv_1 toward PRE and PAR, SUBdd_1 toward medial RSPv, and SUBdv_1 toward lateral RSPv, which substantially revises the earlier view of O’Mara et al. (2001) that PRE and PAR receive only weak SUB input. ENT projections show complementary banding with SUBdd_1 targeting lateral ENT and SUBdv_1 targeting intermediate ENT, while Layer 4 ENT projections diverge substantially between regions, with SUBdv_4 (100.0%) exceeding SUBdd_4 (60.0%).

Layer 4 neurons confirm their CT-class identity through preferentially targeting anterior thalamic nuclei, LD, and RE, which are nuclei strongly implicated in head direction coding, spatial memory, and retrosplenial circuit coordination (Dillingham & Vann, 2019), with SUBdd_4 biased toward more caudal portions of these nuclei and SUBdv_4 more rostrally.

The most notable finding from the SUBdd/dv dataset is the identification of area prostriata (APR) as a collateral target of Layer 1 neurons. APR is a multimodal limbic structure that receives projections from visual and auditory cortices, subiculum, presubiculum, and anterior thalamic nuclei, and plays a critical role in fast processing of peripheral visual motion (Hu et al., 2020; Mikellidou et al., 2017). APR projections were found in 22.6% of SUBdd_1 and 26.6% of SUBdv_1 neurons, and importantly, APR projections exclusively co-occurred with subicular complex collaterals. Specifically, over 50% of neurons projecting to POST, PRE, and PAR, also had an APR collateral in both SUBdd_1 and SUBdv_1. Because APR in turn projects back to POST and PRE (Chen et al., 2021), Layer 1 neurons appear to participate in a reciprocal visuospatial circuit linking the subicular complex and APR that has not been previously characterized at single-cell resolution. Together, these data support the long-standing view of the distal SUB as a structured interface between hippocampus and dispersed visuospatial navigation networks (Sun et al., 2024), while revealing a higher degree of single-cell collateralization and topographic specificity than previously appreciated.

### d. Proubiculum (ProSUB)

Population-level tracing and gene expression analysis from the HGEA established that the ProSUB is primarily dominated by septo-hypothalamic outputs, including LS, hypothalamus, ACB, and MBO, and is distinguished from SUBdd/dv by near-absent RSPv connectivity, indicating a fundamental shift in cortical output profile (Bienkowski et al., 2018). Layer 3 neurons were classified as ET-class and Layer 4 as CT-class with RE as the dominant thalamic target, and earlier tracing studies suggested proximal SUB projections to prefrontal cortex, though primarily attributed to ventral rather than dorsal SUB regions (Jay & Witter, 1991; Finch, 1993; O’Mara et al., 2001). The MPA single-neuron data confirm these findings at single-cell resolution, where the LS (95.5%), hypothalamus (90.1%), ACB (81.1%), and MBO (79.3%) dominance is seen across both layers with near absent RSPv connectivity (1.8%), validating the cortical output shift as a genuine cell-type organizational feature rather than a sampling artifact. At the laminar level, ET-class Layer 3 shows greater prevalence of PFC (35.6% vs. 17.2%), amygdala, and PERI/ECT projections relative to CT-class Layer 4, consistent with an ET-class output profile targeting both cortical and subcortical limbic structures. ProSUB Layer 4 targets RE at the highest rate of any SUB Layer 4 population (75.0%), with a thalamic profile that shifts away from the anterior visuospatial thalamic nuclei that dominates SUBdd/dv Layer 4 toward midline and intralaminar nuclei. This intermediate thalamic profile, connecting between the dominant anterior thalamic-projecting pattern of SUBdd/dv_4 and the midline thalamic-projecting pattern of SUBv/SUBvv_4, is consistent with ProSUB’s structural and positional transition between these two major subdivisions.

Beyond supporting established connectivity patterns, the MPA single-cell data reveals several previously uncharacterized features of ProSUB output. Most notable is the high rate of non-canonical back projections to CA1 (Sun et al., 2019), with 60.4% of ProSUB sending collaterals to dorsal CA1 and 19.5% to CA1i/v. This rate far exceeds that observed in SUBdd/dv, suggesting that the ProSUB plays a specialized role in hippocampal circuit regulation. SUB projections to midbrain structures were also identified, a finding that is discussed in detail in the midbrain section below. Finally, while ENT projections are well represented in the ProSUB, the topographic organization of these projections differs from the SUBdd/dv, with ProSUB axons targeting deeper layers of the lateral ENT and distal SUB axons targeting the caudal dorsomedial ENT. This distinction is functionally significant, as the caudal dorsomedial ENT contains the highest density of large-scale grid cells while the lateral ENT is more closely associated with olfactory and object-related information processing, suggesting that ProSUB and SUBdd/dv make fundamentally different contributions to entorhinal-hippocampal circuit function despite both projecting to ENT (Brun et al., 2008).

### e. Ventral Subiculum (SUBv/SUBvv)

Classical anatomical tracing studies established the output profile of the ventral SUB as a primary hippocampal output to septo-hypothalamic, limbic, and striatal circuits, with dense projections to the LS, hypothalamus, ACB and MBO via the medial corticohypothalamic tract (Canteras & Swanson, 1992; Witter & Groenewegen, 1990). ACB and amygdala also arise preferentially from the ventral SUB (Aylward & Totterdell, 1993; Canteras & Swanson, 1992), and the HGEA later identified a laminar organization underlying these outputs with Layer 2, 3, and 4 neurons showing distinct projection profiles that place SUBv/vv within a brain-wide network that mediates social behavior and neuroendocrine regulation (Bienkowski et al., 2018). The MPA single-neuron reconstructions confirm these findings at single-cell resolution: septo-hypothalamic dominance, IT-class Layer 2 with AON projections, ET-class Layer 3 amygdala projections, and CT-class Layer 4 RE dominance with PVT and PT as additional strong targets are all validated across SUBv and SUBvv.

Several features of SUBv/SUBvv connectivity emerge for the first time from the MPA single-cell data. Most notably, 52.5% of SUBv_2 IT-class neurons extended AON collaterals into the prefrontal cortex, establishing a direct SUB-to-prefrontal pathway that has not been previously described at single-cell resolution. While PFC projections from the ventral SUB have been attributed to ventral SUB populations, the single-cell data indicates that this pathway is specifically enriched in Layer 2 neurons. Within the thalamus, SUBvv_3 ET-class neurons show a pronounced PVT bias (50.0%) compared to SUBv_3 (26.1%), suggesting specialized midline thalamic routing in the most ventral tip of the SUB that may reflect SUBvv’s differential connectivity with endocrine-regulating hypothalamic pathways. Consistent with this, SUBvv shows stronger connectivity with sexually-dimorphic brain areas and medial hypothalamic regions compared to SUBv, which more closely resembles ProSUB in its output topography (Bienkowski et al., 2018; Miller & Aoto, 2025). Finally, projections to PAG and SC were identified in 46.3% of SUBv_4 and 10.0% of SUBvv_4 CT-class neurons via ascending axonal branches that collateralize through the PVT, a finding discussed in detail in the midbrain section below.

### f. SUB projections to Midbrain

Among the most notable cross-regional findings emerging from the ProSUB and SUBv/SUBvv analyses is a set of descending midbrain projections, specifically the PAG, precomissural nucleus (PRC), and SC, a pathway that has received little attention since first described by Köhler (1990). Re-examination of Mouse Connectome Project PHA-L anterograde tracing confirms this projection, although AAV viral tracing with less than 3-4 weeks may not sufficiently label this collateral that is roughly 20mm in axonal path length from the soma (Bienkowski et al., 2018)http://www.mouseconnectome.org (Bienkowski et al., 2018). The MPA data, which uses AAV injections in mice with longer survival times, reveals that these projections arise mostly from ProSUB (ProSUB_3 = 45.8%, ProSUB_4 = 40.4%) and SUBv (SUBv_3 = 32.6%, SUBv_4 = 46.3%), with additional contribution from only 10% of SUBvv_4 neurons. Notably, these axons reach their midbrain targets via two routes: 1) via an ascending branch that first collateralizes near the MBO, 2) caudal extension of the axon that runs entirely through the PVT. Simultaneously engaging midline thalamic, hypothalamic, and midbrain circuits, the functional significance of this pathway is underappreciated given that PAG is a central hub for defensive behavior, stress responses, and fear processing circuits that overlap with the limbic outputs of ProSUB and SUBv (Bandler & Shipley, 1994; Benarroch, 2012). Meanwhile, the SC integrates multisensory information and generates orienting and defensive responses to salient stimuli (Evans et al., 2018; Wei et al., 2015), and its receipt of subicular input suggests a role for the ProSUB and SUBv in directing behavioral responses toward salient environmental stimuli with either positive or negative valence. Together, these projections position the ProSUB and SUBv as regions essential for not only limbic and emotional regulation but also for descending control of allostatic behaviors and warrant further investigation.

## V: Conclusion

### a. From Single Cells to Brain-Wide Networks

This study provides a comprehensive, single-cell validation of the HGEA’s laminar and columnar organization of the SUB. Neurons classified into 12 HGEA cell-type groups relatively reproduced the expected 3D spatial boundaries, confirming that the gene expression-based divisions correspond to functionally distinct projection populations. Key findings include: more than 80% of dorsal SUB Layer 1 neurons co-project to both RSPv and MBO, substantially exceeding prior dual-retrograde estimates; ProSUB and SUBv send previously underappreciated descending projections to the PAG and SC via routes through the PVT and MBO; and IT-class SUBv Layer 2 neurons establish a direct pathway to the prefrontal cortex that has not been described at single-cell resolution. Together, these data reveal that axon collateralization in the SUB is not random but instead a functional feature, with individual neurons conforming to layer-specific motifs while contributing structural heterogeneity across the population. This dataset provides the cell-type-specific wiring detail needed for the next generation of multiscale computational models to accurately simulate how hippocampal output coordinates activity across the brain’s spatial, limbic, and neuroendocrine networks. Incorporating realistic wiring diversity, like that across HGEA-defined cell types, rather than relying on homogenous neuron populations, will be essential for understanding how distributed brain networks give rise to complex behavior.

## Abbreviations

### SUB Regions

SUBdd: Dorsal-dorsal subiculum
SUBdv: Dorsal-ventral subiculum
SUBv: Ventral subiculum
SUBvv: Most ventral subiculum
ProSUB: Prosubiculum

### Frameworks & Atlases

HGEA: Hippocampus Gene Expression Atlas
MPA: Mouse Projectome Atlas C
CFv3: Allen Common Coordinate Framework version 3

### Cell Type Classes

IT: Intratelencephalic neuron
ET: Extratelencephalic neuron
CT: Corticothalamic neuron

### Subicular Complex & Parahippocampal

POST: Postsubiculum
PRE: Presubiculum
PAR: Parasubiculum
APR: Area prostriata
ENT: Entorhinal cortex
PERI/ECT: Perirhinal cortex / Ectorhinal cortex

### Cortical Regions

RSPv: Retrosplenial cortex, ventral part
PFC: Prefrontal cortex
IL: Infralimbic cortex
PL: Prelimbic cortex
TT: Taenia tecta
ORB: Orbital cortex
AON: Anterior olfactory nucleus

### Amygdala

BLA/BMA: Basolateral amygdala / Basomedial amygdala
CEA/MEA: Central amygdala / Medial amygdala

### Striatum & Basal Forebrain

ACB: Nucleus accumbens
LS: Lateral septum
BST: Bed nucleus of the stria terminalis

### Hypothalamus & Brainstem

HYP: Hypothalamus
MBO: Mammillary body
PAG: Periaqueductal gray
SC: Superior colliculus

### Thalamus

RE: Nucleus reuniens
AM: Anteromedial thalamic nucleus
AV: Anteroventral thalamic nucleus
AD: Anterodorsal thalamic nucleus
LD: Laterodorsal thalamic nucleus
IAM: Interanteromedial nucleus
PVT: Paraventricular nucleus of the thalamus
PT: Paratenial nucleus

### Hippocampus

SUB: Subiculum
CA1dc: Cornu Ammonis area 1, dorsal-caudal
CA1i/v: Cornu Ammonis area 1, intermediate / ventral
HATA: Hippocampo-amygdalar transition area

### Techniques

fMOST: Fluorescence micro-optical sectioning tomography

## Acknowledgements

This work was supported by the NIH/NIA K01AG066847 (M.S.B), NIH/NIA R01AG092662 (M.S.B), NIH/NIA P01AG052350 (M.S.B), and National Science Foundation Grant 2121164 (M.S.B.).

## References

Aggleton, J. P., & Christiansen, K. (2015). The subiculum. In Progress in Brain Research (Vol. 219, pp. 65–82). Elsevier. 10.1016/bs.pbr.2015.03.003

Aylward, R. L., & Totterdell, S. (1993). Neurons in the ventral subiculum, amygdala and entorhinal cortex which project to the nucleus accumbens: Their input from somatostatin-immunoreactive boutons. Journal of Chemical Neuroanatomy, 6(1), 31–42. 10.1016/0891-0618(93)90005-o

Bandler, R., & Shipley, M. T. (1994). Columnar organization in the midbrain periaqueductal gray: Modules for emotional expression? Trends in Neurosciences, 17(9), 379–389. 10.1016/0166-2236(94)90047-7

Bastos, A. M., Usrey, W. M., Adams, R. A., Mangun, G. R., Fries, P., & Friston, K. J. (2012). Canonical microcircuits for predictive coding. Neuron, 76(4), 695–711. 10.1016/j.neuron.2012.10.038

Benarroch, E. E. (2012). Periaqueductal gray: An interface for behavioral control. Neurology, 78(3), 210–217. 10.1212/WNL.0b013e31823fcdee

Bienkowski, M. S., Bowman, I., Song, M. Y., Gou, L., Ard, T., Cotter, K., Zhu, M., Benavidez, N. L., Yamashita, S., Abu-Jaber, J., Azam, S., Lo, D., Foster, N. N., Hintiryan, H., & Dong, H.-W. (2018). Integration of gene expression and brain-wide connectivity reveals the multiscale organization of mouse hippocampal networks. Nature Neuroscience, 21(11), 1628–1643. 10.1038/s41593-018-0241-y

Bienkowski, M. S., Sepehrband, F., Kurniawan, N. D., Stanis, J., Korobkova, L., Khanjani, N., Clark, K., Hintiryan, H., Miller, C. A., & Dong, H.-W. (2021). Homologous laminar organization of the mouse and human subiculum. Scientific Reports, 11(1), 3729. 10.1038/s41598-021-81362-w

Boccara, C. N., Kjonigsen, L. J., Hammer, I. M., Bjaalie, J. G., Leergaard, T. B., & Witter, M. P. (2015). A three-plane architectonic atlas of the rat hippocampal region. Hippocampus, 25(7), 838–857. 10.1002/hipo.22407

Boccara, C. N., Sargolini, F., Thoresen, V. H., Solstad, T., Witter, M. P., Moser, E. I., & Moser, M.-B. (2010). Grid cells in pre- and parasubiculum. Nature Neuroscience, 13(8), 987–994. 10.1038/nn.2602

Brun, V. H., Solstad, T., Kjelstrup, K. B., Fyhn, M., Witter, M. P., Moser, E. I., & Moser, M.-B. (2008). Progressive increase in grid scale from dorsal to ventral medial entorhinal cortex. Hippocampus, 18(12), 1200–1212. 10.1002/hipo.20504

Campbell, R. E., Zhang, M. Y., Merryweather, D. N., & Cembrowski, M. S. (2026). Divergent hippocampal output via covariant local and long-range neuronal structure. Progress in Neurobiology, 259, 102893. 10.1016/j.pneurobio.2026.102893

Canteras, N. S., & Swanson, L. W. (1992). Projections of the ventral subiculum to the amygdala, septum, and hypothalamus: A PHAL anterograde tract-tracing study in the rat. Journal of Comparative Neurology, 324(2), 180–194. 10.1002/cne.903240204

Cembrowski, M. S., Phillips, M. G., DiLisio, S. F., Shields, B. C., Winnubst, J., Chandrashekar, J., Bas, E., & Spruston, N. (2018). Dissociable Structural and Functional Hippocampal Outputs via Distinct Subiculum Cell Classes. Cell, 173(5), 1280–1292.e18. 10.1016/j.cell.2018.03.031

Cembrowski, M. S., Wang, L., Lemire, A. L., Copeland, M., DiLisio, S. F., Clements, J., & Spruston, N. (2018). The subiculum is a patchwork of discrete subregions. eLife, 7, e37701. 10.7554/eLife.37701

Chen, C.-H., Hu, J.-M., Zhang, S.-Y., Xiang, X.-J., Chen, S.-Q., & Ding, S.-L. (2021). Rodent Area Prostriata Converges Multimodal Hierarchical Inputs and Projects to the Structures Important for Visuomotor Behaviors. Frontiers in Neuroscience, 15. 10.3389/fnins.2021.772016

DeFelipe, J., & Fariñas, I. (1992). The pyramidal neuron of the cerebral cortex: Morphological and chemical characteristics of the synaptic inputs. Progress in Neurobiology, 39(6), 563–607. 10.1016/0301-0082(92)90015-7

Dillingham, C. M., & Vann, S. D. (2019). Why Isn’t the Head Direction System Necessary for Direction? Lessons From the Lateral Mammillary Nuclei. Frontiers in Neural Circuits, 13. 10.3389/fncir.2019.00060

Ding, S.-L., Yao, Z., Hirokawa, K. E., Nguyen, T. N., Graybuck, L. T., Fong, O., Bohn, P., Ngo, K., Smith, K. A., Koch, C., Phillips, J. W., Lein, E. S., Harris, J. A., Tasic, B., & Zeng, H. (2020). Distinct Transcriptomic Cell Types and Neural Circuits of the Subiculum and Prosubiculum along the Dorsal-Ventral Axis. Cell Reports, 31(7), 107648. 10.1016/j.celrep.2020.107648

Evans, D. A., Stempel, A. V., Vale, R., Ruehle, S., Lefler, Y., & Branco, T. (2018). A synaptic threshold mechanism for computing escape decisions. Nature, 558(7711), 590–594. 10.1038/s41586-018-0244-6

Fanselow, M. S., & Dong, H.-W. (2010). Are The Dorsal and Ventral Hippocampus functionally distinct structures? Neuron, 65(1), 7. 10.1016/j.neuron.2009.11.031

Finch, D. M. (1993). Hippocampal, Subicular, and Entorhinal Afferents and Synaptic Integration in Rodent Cingulate Cortex. In B. A. Vogt & M. Gabriel (Eds.), Neurobiology of Cingulate Cortex and Limbic Thalamus: A Comprehensive Handbook (pp. 224–248). Birkhäuser. 10.1007/978-1-4899-6704-6_8

Gao, P., Cao, W., Nitz, D. A., & Xu, X. (2026). Non-Canonical Subiculum Circuit Organization and Function. Hippocampus, 36(2), e70087. 10.1002/hipo.70087

García-Cabezas, M. Á., Zikopoulos, B., & Barbas, H. (2019). The Structural Model: A theory linking connections, plasticity, pathology, development and evolution of the cerebral cortex. Brain Structure & Function, 224(3), 985–1008. 10.1007/s00429-019-01841-9

Hafting, T., Fyhn, M., Molden, S., Moser, M.-B., & Moser, E. I. (2005). Microstructure of a spatial map in the entorhinal cortex. Nature, 436(7052), 801–806. 10.1038/nature03721

Harris, J. A., Mihalas, S., Hirokawa, K. E., Whitesell, J. D., Choi, H., Bernard, A., Bohn, P., Caldejon, S., Casal, L., Cho, A., Feiner, A., Feng, D., Gaudreault, N., Gerfen, C. R., Graddis, N., Groblewski, P. A., Henry, A. M., Ho, A., Howard, R., … Zeng, H. (2019). Hierarchical organization of cortical and thalamic connectivity. Nature, 575(7781), 195–202. 10.1038/s41586-019-1716-z

Hu, J.-M., Chen, C.-H., Chen, S.-Q., & Ding, S.-L. (2020). Afferent Projections to Area Prostriata of the Mouse. Frontiers in Neuroanatomy, 14, 605021. 10.3389/fnana.2020.605021

Ishizuka, N. (2001). Laminar organization of the pyramidal cell layer of the subiculum in the rat. Journal of Comparative Neurology, 435(1), 89–110. 10.1002/cne.1195

Jay, T. M., & Witter, M. P. (1991). Distribution of hippocampal CA1 and subicular efferents in the prefrontal cortex of the rat studied by means of anterograde transport of *Phaseolus vulgaris* -leucoagglutinin. Journal of Comparative Neurology, 313(4), 574–586. 10.1002/cne.903130404

Kinnavane, L., Vann, S. D., Nelson, A. J. D., O’Mara, S. M., & Aggleton, J. P. (2018). Collateral Projections Innervate the Mammillary Bodies and Retrosplenial Cortex: A New Category of Hippocampal Cells. eNeuro, 5(1). 10.1523/ENEURO.0383-17.2018

Kishi, T., Tsumori, T., Ono, K., Yokota, S., Ishino, H., & Yasui, Y. (2000). Topographical organization of projections from the subiculum to the hypothalamus in the rat. The Journal of Comparative Neurology, 419(2), 205–222. 10.1002/(SICI)1096-9861(20000403)419:2%253C205::AID-CNE5%253E3.0.CO;2-0

Kohler, C. (1990). Subicular projections to the hypothalamus and brainstem: Some novel aspects revealed in the rat by the anterograde Phaseotus vulgaris leukoagglutinin (PHA-L) tracing method.

Lein, E. S., Hawrylycz, M. J., Ao, N., Ayres, M., Bensinger, A., Bernard, A., Boe, A. F., Boguski, M. S., Brockway, K. S., Byrnes, E. J., Chen, L., Chen, L., Chen, T.-M., Chi Chin, M., Chong, J., Crook, B. E., Czaplinska, A., Dang, C. N., Datta, S., … Jones, A. R. (2007). Genome-wide atlas of gene expression in the adult mouse brain. Nature, 445(7124), 168–176. 10.1038/nature05453

Lorente De Nó, R. (1934). Studies on the structure of the cerebral cortex. II. Continuation of the study of the ammonic system. Journal Für Psychologie Und Neurologie, 46, 113–177.

Lu, W., Chen, S., Chen, X., Hu, J., Xuan, A., & Ding, S.-L. (2020). Localization of area prostriata and its connections with primary visual cortex in rodent. The Journal of Comparative Neurology, 528(3), 389–406. 10.1002/cne.24760

Mao, X., & Staiger, J. F. (2024). Multimodal cortical neuronal cell type classification. Pflugers Archiv, 476(5), 721–733. 10.1007/s00424-024-02923-2

Mikellidou, K., Kurzawski, J. W., Frijia, F., Montanaro, D., Greco, V., Burr, D. C., & Morrone, M. C. (2017). Area Prostriata in the Human Brain. Current Biology, 27(19), 3056–3060.e3. 10.1016/j.cub.2017.08.065

Miller, C. N., & Aoto, J. (2025). Sex-differences in the ventral subiculum’s microcircuitry and function. Hippocampus.

Moser, M.-B., & Moser, E. I. (1998). Functional differentiation in the hippocampus. Hippocampus, 8(6), 608–619. 10.1002/(SICI)1098-1063(1998)8:6%253C608::AID-HIPO3%253E3.0.CO; 2-7

O’Mara, S. M., Commins, S., Anderson, M., & Gigg, J. (2001). The subiculum: A review of form, physiology and function. Progress in Neurobiology, 64(2), 129–155. 10.1016/S0301-0082(00)00054-X

Pachicano, M., Mehta, S., Hurtado, A., Ard, T., Stanis, J., Breningstall, B., & Bienkowski, M. S. (2025). Laminar organization of pyramidal neuron cell types defines distinct CA1 hippocampal subregions. Nature Communications, 16(1), 10604. 10.1038/s41467-025-66613-y

Peyrache, A., Schieferstein, N., & Buzsáki, G. (2017). Transformation of the head-direction signal into a spatial code. Nature Communications, 8(1), 1752. 10.1038/s41467-017-01908-3

Qiu, S., Hu, Y., Huang, Y., Gao, T., Wang, X., Wang, D., Ren, B., Shi, X., Chen, Y., Wang, X., Wang, D., Han, L., Liang, Y., Liu, D., Liu, Q., Deng, L., Chen, Z., Zhan, L., Chen, T., … Xu, C. (2024). Whole-brain spatial organization of hippocampal single-neuron projectomes. Science, 383(6682), eadj9198. 10.1126/science.adj9198

Ramón y Cajal, S. (with University of Illinois Urbana-Champaign). (1909). Histologie du système nerveux de l’homme & des vertébrés. Paris : Maloine. http://archive.org/details/histologiedusyst01ram

Rosenblum, E. W., Williams, E. M., Champion, S. N., Frosch, M. P., & Augustinack, J. C. (2024). The prosubiculum in the human hippocampus: A rostrocaudal, feature-driven, and systematic approach. The Journal of Comparative Neurology, 532(3), e25604. 10.1002/cne.25604

Saleeba, C., Dempsey, B., Le, S., Goodchild, A., & McMullan, S. (2019). A Student’s Guide to Neural Circuit Tracing. Frontiers in Neuroscience, 13. 10.3389/fnins.2019.00897

Simonsen, Ø. W., Czajkowski, R., & Witter, M. P. (2022). Retrosplenial and subicular inputs converge on superficially projecting layer V neurons of medial entorhinal cortex. Brain Structure and Function, 227(8), 2821–2837. 10.1007/s00429-022-02578-8

Strange, B. A., Witter, M. P., Lein, E. S., & Moser, E. I. (2014). Functional organization of the hippocampal longitudinal axis. Nature Reviews Neuroscience, 15(10), 655–669. 10.1038/nrn3785

Sun, Y., Jin, S., Lin, X., Chen, L., Qiao, X., Jiang, L., Zhou, P., Johnston, K. G., Golshani, P., Nie, Q., Holmes, T. C., Nitz, D. A., & Xu, X. (2019). CA1-Projecting Subiculum Neurons Facilitate Object-Place Learning. Nature Neuroscience, 22(11), 1857–1870. 10.1038/s41593-019-0496-y

Sun, Y., Nitz, D. A., Xu, X., & Giocomo, L. M. (2024). Subicular neurons encode concave and convex geometries. Nature, 627(8005), 821–829. 10.1038/s41586-024-07139-z

Swanson, L. W. (1981). Tracing Central Pathways with the Autoradiographic Method,. Journal of Histochemistry & Cytochemistry, 29(1A_suppl), 117–124. 10.1177/29.1A_SUPPL.7288150

Swanson, L. W., & Cowan, W. M. (1975). Hippocampo-Hypothalamic Connections: Origin in Subicular Cortex, Not Ammon’s Horn. Science, 189(4199), 303–304. 10.1126/science.49928

Umaba, R., Kitanishi, T., & Mizuseki, K. (2021). Monosynaptic connection from the subiculum to medial mammillary nucleus neurons projecting to the anterior thalamus and Gudden’s ventral tegmental nucleus. Neuroscience Research, 171, 1–8. 10.1016/j.neures.2021.01.006

Van Groen, T. (2001). Entorhinal cortex of the mouse: Cytoarchitectonical organization. Hippocampus, 11(4), 397–407. 10.1002/hipo.1054

Wei, P., Liu, N., Zhang, Z., Liu, X., Tang, Y., He, X., Wu, B., Zhou, Z., Liu, Y., Li, J., Zhang, Y., Zhou, X., Xu, L., Chen, L., Bi, G., Hu, X., Xu, F., & Wang, L. (2015). Processing of visually evoked innate fear by a non-canonical thalamic pathway. Nature Communications, 6(1), 6756. 10.1038/ncomms7756

Winnubst, J., Bas, E., Ferreira, T. A., Wu, Z., Economo, M. N., Edson, P., Arthur, B. J., Bruns, C., Rokicki, K., Schauder, D., Olbris, D. J., Murphy, S. D., Ackerman, D. G., Arshadi, C., Baldwin, P., Blake, R., Elsayed, A., Hasan, M., Ramirez, D., … Chandrashekar, J. (2019). Reconstruction of 1,000 projection neurons reveals new cell types and organization of long-range connectivity in the mouse brain. Cell, 179(1), 268–281.e13. 10.1016/j.cell.2019.07.042

Witter, M. P. (2006). Connections of the subiculum of the rat: Topography in relation to columnar and laminar organization. Behavioural Brain Research, 174(2), 251–264. 10.1016/j.bbr.2006.06.022

Witter, M. P., & Groenewegen, H. J. (1990). The subiculum: Cytoarchitectonically a simple structure, but hodologically complex. Progress in Brain Research, 83, 47–58. 10.1016/s0079-6123(08)61240-6

Woych, J., Ortega Gurrola, A., Deryckere, A., Jaeger, E. C. B., Gumnit, E., Merello, G., Gu, J., Joven Araus, A., Leigh, N. D., Yun, M., Simon, A., & Tosches, M. A. (2022). Cell-type profiling in salamanders identifies innovations in vertebrate forebrain evolution. Science, 377(6610), eabp9186. 10.1126/science.abp9186

Xu, X., Sun, Y., Holmes, T. C., & López, A. J. (2016). Noncanonical connections between the subiculum and hippocampal CA1. Journal of Comparative Neurology, 524(17), 3666–3673. 10.1002/cne.24024

Yao, Z., van Velthoven, C. T. J., Kunst, M., Zhang, M., McMillen, D., Lee, C., Jung, W., Goldy, J., Abdelhak, A., Aitken, M., Baker, K., Baker, P., Barkan, E., Bertagnolli, D., Bhandiwad, A., Bielstein, C., Bishwakarma, P., Campos, J., Carey, D., … Zeng, H. (2023). A high-resolution transcriptomic and spatial atlas of cell types in the whole mouse brain. Nature, 624(7991), 317–332. 10.1038/s41586-023-06812-z

Zhang, J., Yao, M., Jiang, T., Li, A., Gong, H., He, M., Luo, Q., & Li, X. (2025). A dorsal subiculum-medial mammillary body pathway for spatial memory. Molecular Psychiatry, 30(11), 5045–5057. 10.1038/s41380-025-03087-w

Zhang, S.-Y., Chen, S.-Q., Zhang, J.-Y., Chen, C.-H., Xiang, X.-J., Cai, H.-R., & Ding, S.-L. (2022). The effects of bilateral prostriata lesions on spatial learning and memory in the rat. Frontiers in Behavioral Neuroscience, 16. 10.3389/fnbeh.2022.1010321

